# Variability and correlations among vital rates and their influence on population growth in mule and black-tailed deer

**DOI:** 10.1101/2024.03.15.585316

**Authors:** Joel Ruprecht, Tavis D. Forrester, Darren A. Clark, Michael J. Wisdom, Joshua B. Smith, Taal Levi

**Affiliations:** Oregon State University, 104 Nash Hall, Corvallis, OR 97331, USA; 800 East Beckwith Avenue, Rocky Mountain Research Station, USDA Forest Service, Missoula, MT 59801, USA; Oregon Department of Fish and Wildlife, 1401 Gekeler Lane, La Grande, OR 97850, USA; Pacific Northwest Research Station, USDA Forest Service, 1401 Gekeler Lane, La Grande, OR 97850, USA; Oregon Department of Fish and Wildlife, 107 20^th^ Street, La Grande, OR 97850

**Keywords:** large herbivore, life-stage simulation analysis, matrix model, *Odocoileus hemionus*, survival, ungulate

## Abstract

To reverse observed range-wide population declines, managers of mule and black- tailed deer (*Odocoileus hemionus*) require information on the vital rates and life stages that are most influential to population growth for which to target management actions. We conducted a range-wide literature review and used hierarchical models to provide biological descriptions of mule and black-tailed deer vital rates, their variability, and how they correlate with one another. We then used matrix models and life-stage simulation analysis to determine the individual vital rates that contributed most to lambda, i.e., annual population growth rate. Adult female survival was the vital rate with the greatest ability to predict lambda and explained 62% of the variation in population growth. While annual juvenile survival explained 44% of the variation in lambda, summer and winter juvenile survival by themselves were far less explanatory than adult female survival. Winter fawn:doe ratios, a metric often collected by management agencies, only explained 10% of the variation in lambda. When adult female survival was 0.84, our simulations estimated an equal probability that a population would increase versus decrease, and correspondingly, the estimate of lambda was 1.0 with a 95% credible interval of 0.88–1.14. Simulations suggested that populations with adult survival rates less than 70% would always decline, but as survival increased beyond this value lambda increased linearly and never plateaued. In contrast, the probability of observing a stable lambda plateaued when annual juvenile survival reached approximately 0.5. The mean lambda calculated from all simulated values within the observed range of vital rate values across the species’ geographical distribution was 0.975, and in 61% of the simulations lambda was < 1. After 20 years, we estimated that this distribution of lambda values would cause populations to decrease in 92% of instances with a mean decrease of 44%. Our results align with the observed declines in mule deer populations throughout their range over recent decades and suggest that these trends will continue until management can improve survival of adult females.

---

The application of matrix models to quantify animal population growth, and the associated perturbation analysis to understand the “importance” of individual vital rates to population growth, has underpinned many applications of wildlife population ecology and management in the last century. Lewis (1942) and Leslie (1945) first demonstrated that matrices could be populated with age-specific vital rates and that the dominant eigenvalue was equivalent to the long-term geometric growth rate (hereafter, *lambda* or λ). Lefkovitch (1965) generalized the age-based (“Leslie”) matrix such that vital rates could be stage-based, and these “Lefkovitch” matrices have been important for organisms whose vital rates can easily be grouped into stage classes instead of the more restrictive data requirements of age-specific rates. Caswell (1978) was the first to apply perturbation analysis (i.e. sensitivity or elasticity analysis) to the dominant eigenvalue, demonstrating that different vital rates or different age or stage classes had variable contributions to lambda, with population growth being highly sensitive to certain vital rates but nearly invariant to others. For applied wildlife ecology, this meant that alternative management or conservation strategies could be evaluated with the expected overall effect on population growth rate quantified *a priori*. Perhaps the most classic example of this was the determination that turtle excluder devices, by protecting the age class with the highest sensitivity (large juveniles), would yield the greatest increase in population size when evaluating several management alternatives for loggerhead turtles (*Caretta caretta*) under a limited budget and high public scrutiny (Crowder et al. 1994).

While perturbation analysis is a powerful tool in applied wildlife management, it provides an incomplete understanding of how vital rates ultimately influence population growth. This is because sensitivities and elasticities only quantify how lambda would be affected *if* vital rates changed, but they tell nothing about *how* they change, especially in the context of real systems. This has important implications when evaluating management alternatives, because it would not be efficient to manage for the vital rate with the highest elasticity if that vital rate has almost no potential to vary in nature or in response to a management action. This is often a common scenario confronting managers seeking to improve population performance because the evolution of life history strategies has resulted in an inverse relationship between elasticity and variability in vital rates (Pfister 1998). Correspondingly, the vital rates with the greatest capacity to influence lambda are buffered the most from environmental variation and thus the least likely to change (Gaillard and Yoccoz 2003). Therefore, some combination of elasticity and variability within vital rates leads to certain ones explaining more variation in lambda than others. Though there is a very nuanced distinction between the elasticity of a vital rate to lambda versus the amount of variation it explains in lambda, the difference is important, and two analytical methods—lifetable response experiments Caswell (1989) and life-stage simulation analysis (LSA; Wisdom et al. (2000))—attempt to reconcile this difference. Lifetable response experiments extend classical perturbation analysis to include variation in vital rates using analytical methods, whereas LSA uses simulations from probability distributions characterizing vital rates to evaluate lambda under a range of hypothetical scenarios to determine the influence of each vital rate on population growth. While both methods are useful, only LSA allows one to make probabilistic statements about population growth based on the influence of each vital rate as it varies with all other rates under defined probability distributions (Wisdom et al. 2000).

Large herbivores provide an ideal case study for employing LSA to understand the nuances between the *potential* ability for vital rates to influence lambda, and the *realized* influence in natural systems occurring either through environmental variation or management action. This is because a strong body of literature supports the paradigm in which their life history strategy employs high and constant adult female survival, but lower and more variable juvenile survival (Gaillard et al. 1998, Gaillard et al. 2000). Perturbation analysis consistently ranks adult female survival as having the highest elasticity (Gaillard and Yoccoz 2003), but in most systems it changes little because of selective forces buffering it from the effects of population density and environmental variation. As such, the prevailing dogma of large herbivore life history theory maintains that most of the interannual variation in population growth rate arises through variation in juvenile survival (Gaillard et al. 2000); however, the few studies to formally quantify this have not always agreed. For example, in a LSA for elk (*Cervus canadensis*) compiled from populations across their North American distribution, Raithel et al. (2007) estimated that 75% of the variation in lambda could be explained by calf survival. However, in a more limited geographical extent in Montana, calf survival explained less than 20% of the variation in lambda and adult female survival explained as much as 80% (Eacker et al. 2017), similar to the LSA results in a congener, the sika deer (*Cervus nippon;* Miura and Tokida (2009)). In woodland caribou (*Rangifer tarandus caribou*), LSA determined that 54% of the variation in lambda was explained by adult survival and 43% by recruitment (DeCesare et al. 2012). In Sitka black-tailed deer (*Odocoileus heminous sitkensis*), lifetable response experiments revealed that adult female survival, by virtue of having almost no process error over the 3 years of a study in southeast Alaska, contributed almost nothing to lambda despite it being the most elastic vital rate (Gilbert et al. 2020).

Mule deer (*Odocoileus hemionus*) and black-tailed deer (*Odocoileus hemionus columbianus*) are species of high management concern in the 21^st^ century due to declines from historic numbers and consequently, many state management agencies across the western United States have recently initiated large-scale demographic studies. As managers seek to reverse declining population trends, they will benefit from a thorough understanding of how stage-specific vital rates ultimately contribute to population growth, and how certain vital rates may or may not be reliable predictors of lambda. While perturbation analyses specific to mule and black-tailed deer have demonstrated that they too display high elasticity to adult female survival (Bishop et al. 2009, Marescot et al. 2015, Gilbert et al. 2020, Lukacs and Nowak 2023), without proper consideration of both elasticity and variability, the degree to which adult female survival ultimately correlates with lambda cannot be fully understood. Local and regional studies have estimated process standard deviation (i.e. biological variation that is separate from sampling variation) surrounding adult female survival to be quite low (0.03–0.07) (Unsworth et al. 1999, Lukacs et al. 2009) which would suggest that canalization has led to constancy and buffering from environmental variation. However, mean range-wide estimates of mule deer adult female survival (Forrester and Wittmer 2013) are lower than those of elk (Raithel et al. 2007), indicating that mule deer may have a greater capacity for adult survival to influence lambda given that a broader range of values could be attained (Sæther and Bakke 2000). Alternatively, mule deer have larger litter sizes and can reproduce earlier than elk, and the greater variation in reproductive output could plausibly cause fecundity to explain more variation in lambda than in elk.

Our objectives were to 1) conduct a range-wide literature review of mule and black-tailed deer vital rates, 2) use hierarchical models and Bayesian meta-analysis methods to characterize vital rates and correlations among vital rates using probability distributions, 3) employ life-stage simulation analysis (LSA) to determine the vital rates that explain the most variation in population growth, and 4) use output from the LSA to estimate the probability a population would increase given the observation of a point estimate of an individual vital rate. We hypothesized that similar to many other large herbivores, population growth would be most influenced by variation in juvenile survival (Gaillard et al. 1998, Gaillard et al. 2000). These results will advance our understanding of how individual vital rates contribute to mule deer population growth across their geographic distribution that can be used to guide management.

## METHODS

We first specified a post-breeding, female-only matrix model using three stages (juveniles [age 0], yearlings [age 1], and adults [age 2+]) to quantify mule deer population growth (Equation 1). Our model is functionally similar to the matrix described by Bishop et al. (2009) and identical to that described by Gilbert et al. (2020). This model assumes that yearling pregnancy, fecundity, and survival rates differ from those of adults, but that all deer greater than 2 years old have the same rates. We did not assume senescence in reproductive or survival rates due to a paucity of information reported for the species. The matrix took the form:

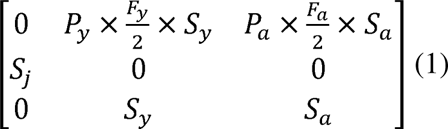

where *P* is the annual pregnancy rate, *F* is fecundity, and *S* is survival, and where *j* subscripts indicate vital rates of juveniles, *y* yearlings, and *a* adults. Fecundity, or the product of litter size (*LS*) and pregnancy, is divided by 2 to assume an equal sex ratio of male and female fawns. We used the product of summer juvenile survival (*S_j sum_*; 0–6 months) and winter survival (*S_j win_*, 7– 12 months) to estimate 12-month juvenile survival, *S_j_*. In the modeling below, litter size is given that (1 – *p*) × 1 + *p* × 2 = *p* + 1, where *p* is twinning rate. This was done so that represented as the proportion of the parturient females that have twins. This derivation works twinning rate could be modeled on the logit scale along with the other vital rates with biological bounds of 0 and 1. After modeling, we added 1 to the twinning rate which converts the value back to number of fawns per pregnant female. This approach assumes that a population’s average litter size could not be > 2 which is reasonable given that mule and black-tailed deer only rarely have triplets (Monteith et al. 2023).

We then conducted a comprehensive literature review to characterize each vital rate in our matrix (Supporting Information S1). We used Bayesian logit-normal hierarchical models to estimate means, variances, and correlations among the vital rates in our matrix (Lukacs et al. 2009). A multivariate logit-normal distribution is the preferred option to jointly model the correlations among every pair of vital rates within a single model in addition to the mean and variance of each individual vital rate. However, this would require that every study reported every vital rate needed for the matrix model, and this was not possible given that the overwhelming majority of studies in our review reported only one or several vital rates. We therefore fit two sets of models—one to characterize the means and variances of each individual vital rate independently, and another set that estimated correlations among each pair of vital rates using subsets of the data for studies that reported multiple vital rates. The first set of models (2, 3) used all point estimates of each vital rate and took the form:

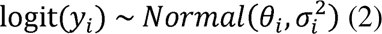

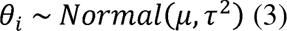

where *y_i_* is the vital rate estimate from study *i*, *θ_i_* is the true mean vital rate in study *i* on the logit scale, 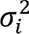 is the within study *i* variance (or *sampling* variance) of the logit-transformed vital rate, *μ* is the true, grand mean of the vital rate on the logit scale across studies, and *τ*^2^ is the among study variance (or *process* variance) of the logit-transformed vital rate. Equation 2 specifies that the logit of each vital rate reported in our literature review (*y_i_*) is normally distributed, with mean *θ_i_* and sampling variance, 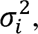 which is derived from the standard error associated with each study’s reported vital rate. Note that 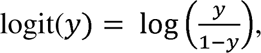 and to express a standard deviation for a variable *y* on the logit scale, one must use 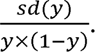 Hence, in our study 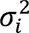 for each study was specified as 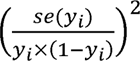 where *se*(*y_i_*) is the reported standard error of each study’s vital rate estimate. Within Equation 3, *μ* and *τ*^2^ provide a rangewide, biological description of the vital rate on the logit scale. Note that the inverse logit can be used to back-transform *μ* from the logit scale to the normal scale using inverse 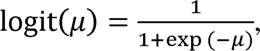 and *τ*^2^ can be back-transformed using *τ*^2^ × (*inverse logit*(*μ*) × (1 - *inverse logit*(*μ*)))^2^. We only report parameter values on their normal, back-transformed scales.

The second set of models (4, 5) used a subset of vital rate observations to estimate correlations among each pair of vital rates. This was necessary because few studies reported all vital rates and thus to estimate correlations we could only use a subset of the studies reporting matched pairs of vital rates. These models were similar to the first set but assumed a bivariate normal distribution allowing for correlations to be estimated:

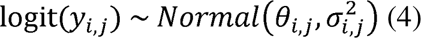

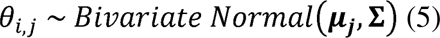

where all parameters are as described above with *i* indexing each study and *j* indexing each pair of vital rates being jointly modeled, and **Σ** is the covariance matrix. Within **Σ**, diagonal entries represent process variance *τ*^2^, and off diagonal elements represent covariances, which for a pair of vital rates is indicated by *τ*_1_ × *τ*_2_ × *ρ*, where *ρ* is the correlation coefficient between the two vital rates on the logit scale.

We used an Inverse Gamma(1, 1) prior for *τ*^2^, a Normal(0, *σ*^2^ = 2) prior for *μ*, and a Normal(0, *σ*^2^ = 4) prior for *ρ*, truncated at -1, 1. We fit models in Nimble using 3 chains of 50,000 iterations after a 50,000 iteration burn-in phase. We ensured 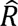 values were less than 1.05 and visually inspected traceplots for convergence.

To conduct life-stage simulation analysis, we generated a random draw from a multivariate logit-normal distribution for each of the 150,000 posterior samples of the means and variances of each vital rate estimated using Equation 3, and with correlations among each pair of vital rates (*ρ*) estimated within Equation 5. This resulted in a dataset of 150,000 values for each of 8 vital rates with the covariance structure as estimated and described above. For each of the 150,000 draws of correlated vital rates, we populated a matrix (Equation 1) and calculated the dominant eigenvalue as the expected population growth rate given that set of vital rate values. We also calculated analytical elasticity values for each vital rate. We then used simple linear regression to model lambda as a function of each individual vital rate, and calculated the coefficient of determination (*R^2^*) to determine the proportion of variance in lambda explained by each vital rate.

We also sought to evaluate how well winter fawn:doe ratios could predict lambda because such metrics of herd composition are easily and often collected by wildlife managers. We therefore calculated what the expected fawn:doe ratio would be during winter for each set of randomly-drawn fecundity and adult and juvenile summer (June-December) survival rates. This was achieved by taking the ratio in which the numerator quantified the number of fawns produced per adult female that survived and could be counted during winter herd composition surveys, and the denominator quantified the proportion of yearling and adult deer that survived until winter and could be counted during winter herd composition surveys. Note that in the numerator, only adult deer produce offspring because yearlings would not yet have had a chance to produce offspring at 1.5 years old, but in the denominator we specify that there were yearling females in addition to the breeding adult females that were present during herd composition surveys and could be counted as adults. This was achieved by using a correction factor of 1.176, because age-specific survival rates from two populations suggested that there should be 0.176 yearlings for every adult female (Bishop et al. 2009, Monteith et al. 2014). Thus,

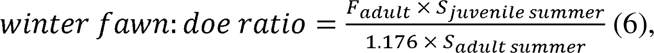

where *P* is pregnancy rate, *F* is fecundity (i.e., the product of pregnancy and litter size), and *S* is survival. This equation required estimates of summer (∼June 1–30 November) adult female survival which we estimated from four sources (Bender et al. 2007, Bishop et al. 2009, Hurley et al. 2011, Monteith et al. 2014). We then predicted a summer adult survival rate value from each annual adult survival rate by raising the latter to the power of *x*, where 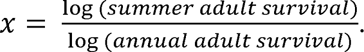 We assumed yearlings and adults had similar summer survival rates so we did not separately calculate a summer survival rates for yearlings.

Next, we used the simulated vital rates to make probabilistic statements about lambda given the observation of a point estimate of either adult survival, juvenile survival, or winter fawn:doe ratios. We did this in two ways. First, across all plausible values of the vital rate of interest, we plotted the proportion of instances in our simulated dataset in which lambda was ≥ 1. This allowed us to visually quantify whether there were nonlinearities in this relationship; for example, a line that plateaued would indicate diminishing returns such that an increase in the vital rate beyond a certain value would not lead to a higher probability of a stable population. Second, we plotted the mean and 2.5% and 97.5% quantiles of lambda across all plausible values of the vital rate of interest. We constructed these plots as a tool to understand the range of possible growth rates given the observation of a single vital rate point estimate.

Finally, we summarized the values of lambda as estimated from 150,000 matrix model simulations and calculated the proportion of instances that resulted in a lambda < 1. We then assessed how this lambda distribution could influence populations over a 20-year time period. To do this, we generated 10,000 population trajectories each beginning with 1,000 individuals (*N*_1_ = 1,000) and iteratively updated *N_t+_*_1_ based on *N_t_* and a randomly drawn value of lambda each year (λ*_t_*) using *N_t+_*_1_ = *N_t_* × λ*_t_*.

## RESULTS

Our literature review yielded 722 annual point estimates (and associated standard errors) across the 8 vital rates needed for our matrix model using 72 different data sources spanning 14 states and 2 provinces (Supporting Information S1, S2). Of the 72 data sources, 60 were conducted on mule deer, 11 on black-tailed deer, and 1 on hybrid mule and black-tailed in their introgression zone. The most commonly-reported vital rates were adult survival and overwinter juvenile survival (Supporting Information S1, S2). The hierarchical logit-normal models fit to data from our literature review provided summaries of each vital rates in terms of its range-wide mean (*x*□) and process standard deviation (*σ*) (Fig. 1, Table 1). Summer (0–6 month) juvenile survival (*x*□ = 0.458, *σ* = 0.146) was lower but less variable than winter (7–12 month) survival (*x*□ = 0.613, *σ* = 0.224). Curiously, when we modeled annual juvenile survival (*x*□ = 0.308, *σ* = 0.130) from studies reporting annual estimates of the vital rate, it was greater than the product of survival estimates of 0–6 month and 7–12 month fawns (Supporting Information S3). Annual yearling (1–2 yrs) female survival (*x*□ = 0.840, *σ* = 0.103) was similar but slightly higher and more variable than for adult (2+ yrs) females (*x*□ = 0.826, *σ* = 0.063). The mean summer adult survival rate was 0.934. Yearling pregnancy (*x*□ = 0.726, *σ* = 0.161) was substantially lower and more variable than adult pregnancy (*x*□ = 0.942, *σ* = 0.030), whereas yearling litter size (*x*□ = 1.345, *σ* = 0.198) was modestly lower than for adults (*x*□ = 1.657, *σ* = 0.138). The most variable vital rates were winter juvenile survival and yearling reproductive rates, whereas those with the highest precision were annual adult survival and adult pregnancy (Fig. 2).

**Figure 1.**
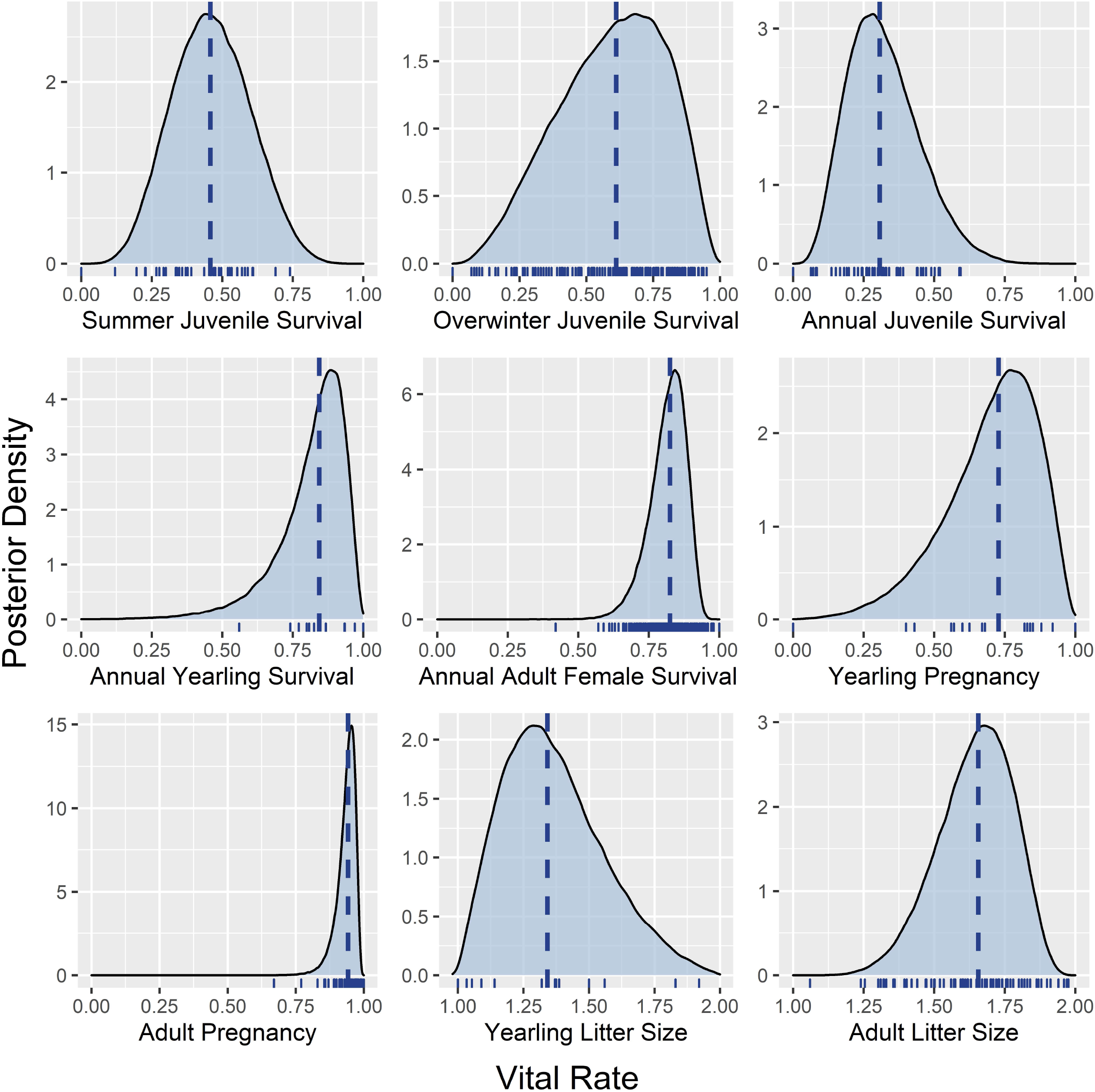
Posterior densities of mule deer vital rates fit using logit-normal hierarchical models summarized from a range-wide literature review. The vertical dotted line displays the posterior mean. The small vertical ticks on the x-axis show mean values of each point estimate used in the meta-analysis. Litter size refers to the number of offspring produced per pregnant female, and thus fecundity (number of offspring per adult female), is the product of pregnancy rate and litter size. Summer juvenile survival is from 0–6 months and winter juvenile survival is from 7–12 months of age.

**Figure 2.**
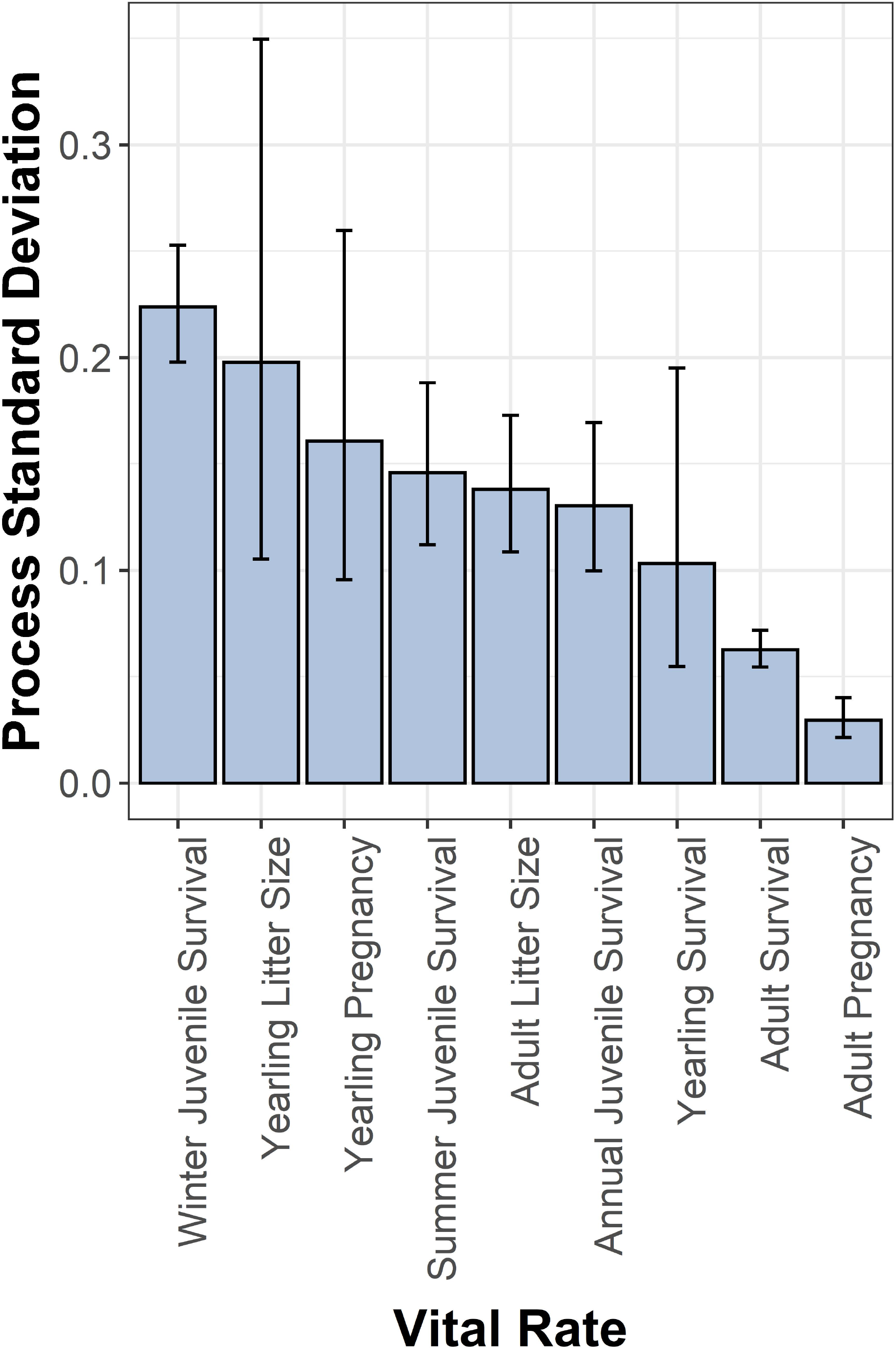
Estimates of process standard deviation (*τ* in Equation 3 on the normal scale) of mule and black-tailed deer vital rates, fit using logit-normal hierarchical models summarized from a range-wide literature review. Error bars represent 95% credible intervals for the estimate of the posterior standard deviation of the vital rate. Note that process standard deviation characterizes biological variability within vital rates and is separate from sampling variance.

**Table 1.** Summary of vital rates used in the LSA. *N* indicates the number of point estimates (annual vital rates). Summer juvenile survival is from 0–6 months, and winter juvenile survival is from 7–12 months. Studies reporting juvenile survival rates over shorter periods (e.g. 0–3 months) were not included.

| **Vital Rate** | **Median** | **Mean** | **Process SD** | ***N*** |
| --- | --- | --- | --- | --- |
| Juvenile Survival (annual) | 0.308 | 0.308 | 0.130 | 47 |
| Juvenile Survival (summer) | 0.458 | 0.458 | 0.146 | 52 |
| Juvenile Survival (winter) | 0.613 | 0.613 | 0.224 | 211 |
| Yearling Survival (annual) | 0.844 | 0.840 | 0.103 | 12 |
| Adult Female Survival (annual) | 0.826 | 0.826 | 0.063 | 249 |
| Yearling Pregnancy | 0.728 | 0.726 | 0.161 | 22 |
| Adult Pregnancy | 0.942 | 0.942 | 0.030 | 78 |
| Yearling Litter Size | 1.342 | 1.345 | 0.198 | 18 |
| Adult Litter Size | 1.657 | 1.657 | 0.138 | 96 |

Correlations (*ρ*) among vital rates could only be estimated for certain vital rates from studies with sufficient matched pairs (see non-NA entries in Fig. 3 and Supporting Information S4: Table S1), and of these, only 7 had Bayesian credible intervals that did not span 0 (colored entries in Fig. 3; bold entries in Supporting Information S4: Table S1). Most surprisingly was the observation that summer and winter juvenile survival were negatively correlated (*ρ* = -0.44). Overwinter juvenile survival was positively correlated with both yearling and adult survival (*ρ* = 0.79 and *ρ* = 0.23, respectively), and adult survival also correlated positively with adult pregnancy (*ρ* = 0.59). Adult litter size was positively correlated with yearling pregnancy (*ρ* = 0.70), adult pregnancy (*ρ* = 0.24), and yearling litter size (*ρ* = 0.74).

**Figure 3.**
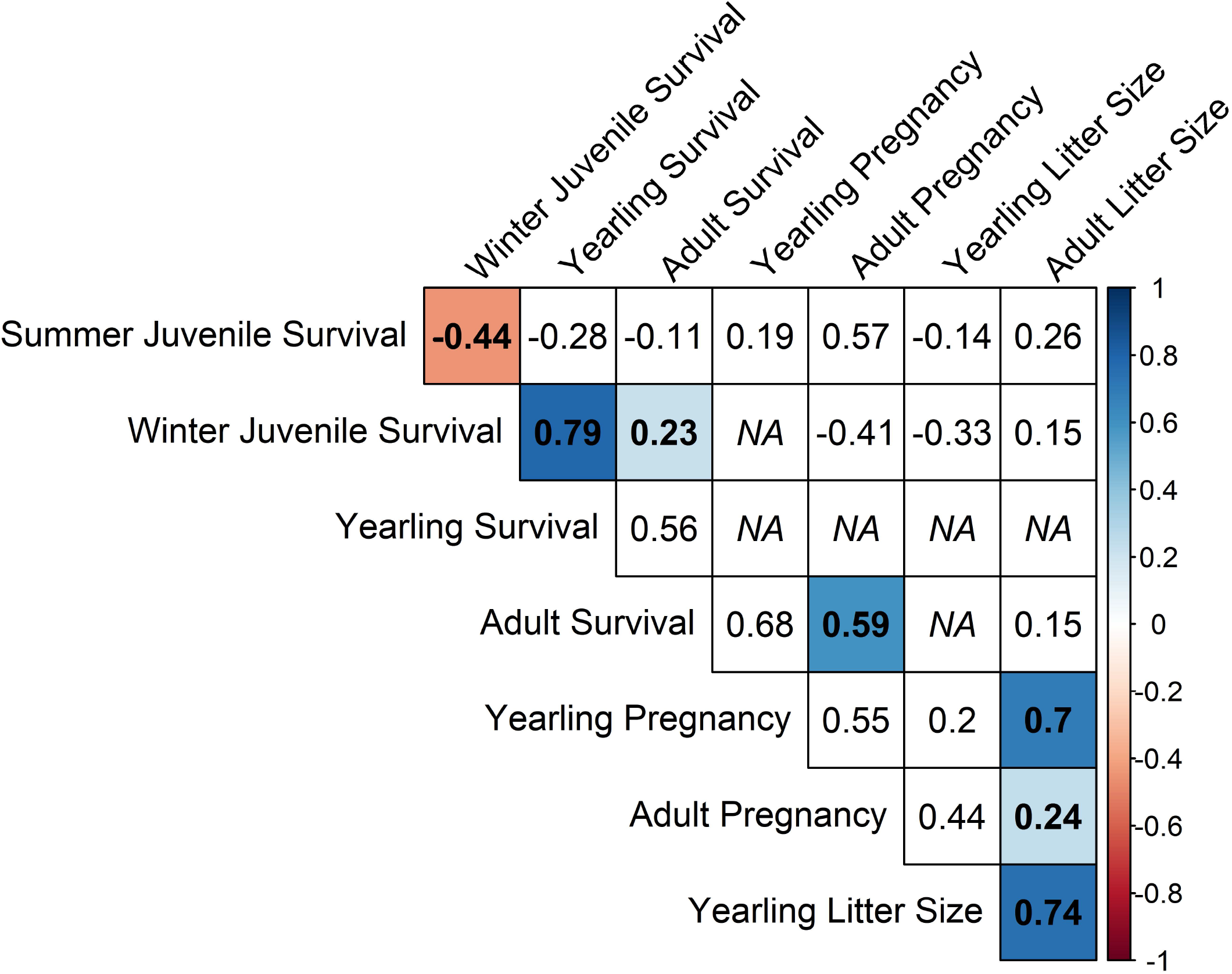
Estimates of the correlations (*ρ*) among pairs of mule and black-tailed deer vital rates fit using bivariate logit-normal hierarchical models summarized from a range-wide literature review. NA entries represent pairs of vital rates with insufficient data to fit models. Colored entries represent correlations whose 95% Bayesian credible intervals do not overlap 0.

Adult female survival was overwhelmingly the vital rate with the highest elasticity value (0.72) and was more than 5 times greater than the next most elastic vital rates at 0.14 (survival of juveniles during summer, overwinter survival of juveniles, and annual survival of yearlings) (Figure 4). The terms constituting adult fecundity (pregnancy and litter size) had elasticity values only slightly lower than juvenile and yearling survival (0.12), and the yearling fecundity terms were the least elastic with values of 0.02.

**Figure 4.**
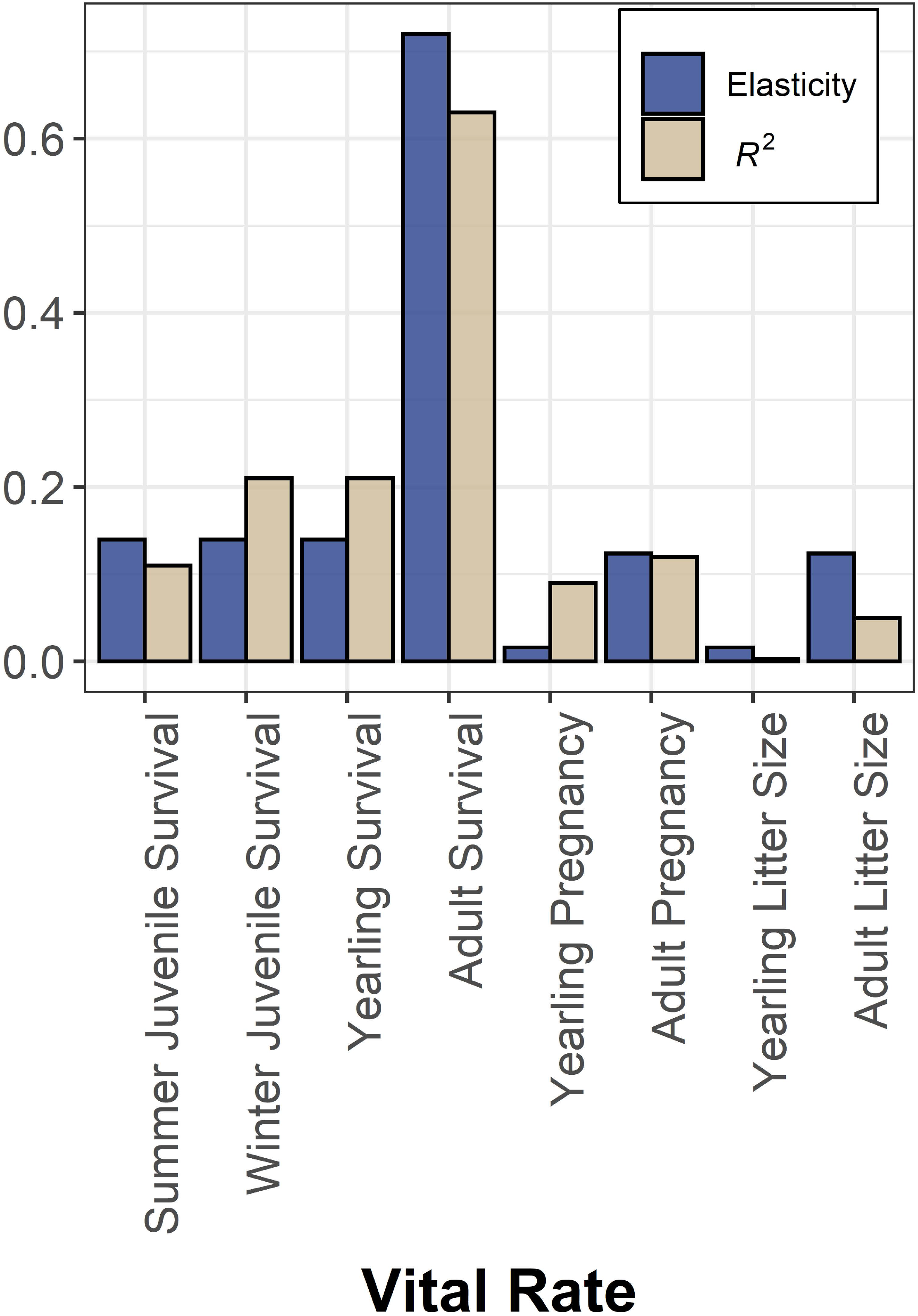
Comparisons of analytical elasticity values from a matrix population model and *R^2^* values from a life-stage simulation analysis in mule and black-tailed deer. Elasticity values measure the magnitude of the change in lambda expected from a proportional change in a vital rate, and thus describe the *potential* ability for vital rates to influence population growth. *R^2^* values (coefficients of determination) in a life-stage simulation analysis measure the amount of variation in lambda that can be explained by individual vital rates under realistic scenarios, and thus describe the *realized* ability for vital rates to influence population growth.

The LSA (Fig. 5) revealed that the majority (62%) of variation in lambda was explained by adult female survival and that summer and winter juvenile survival only explained 10% and 25%, respectively. A separate regression with a single parameter that was the product of summer and winter juvenile survival indicated that annual juvenile survival explained 44% of the variation in lambda. Yearling survival explained 22% of the variation in lambda, and yearling or adult pregnancy and litter size parameters explained <12% of the variation in lambda. When we calculated expected winter fawn:doe ratios using Equation 6, we determined that winter fawn:doe ratios explained only 10% of the variation in lambda. Elasticity values and *R^2^* values from the LSA were similar among vital rates (Fig. 4).

**Figure 5.**
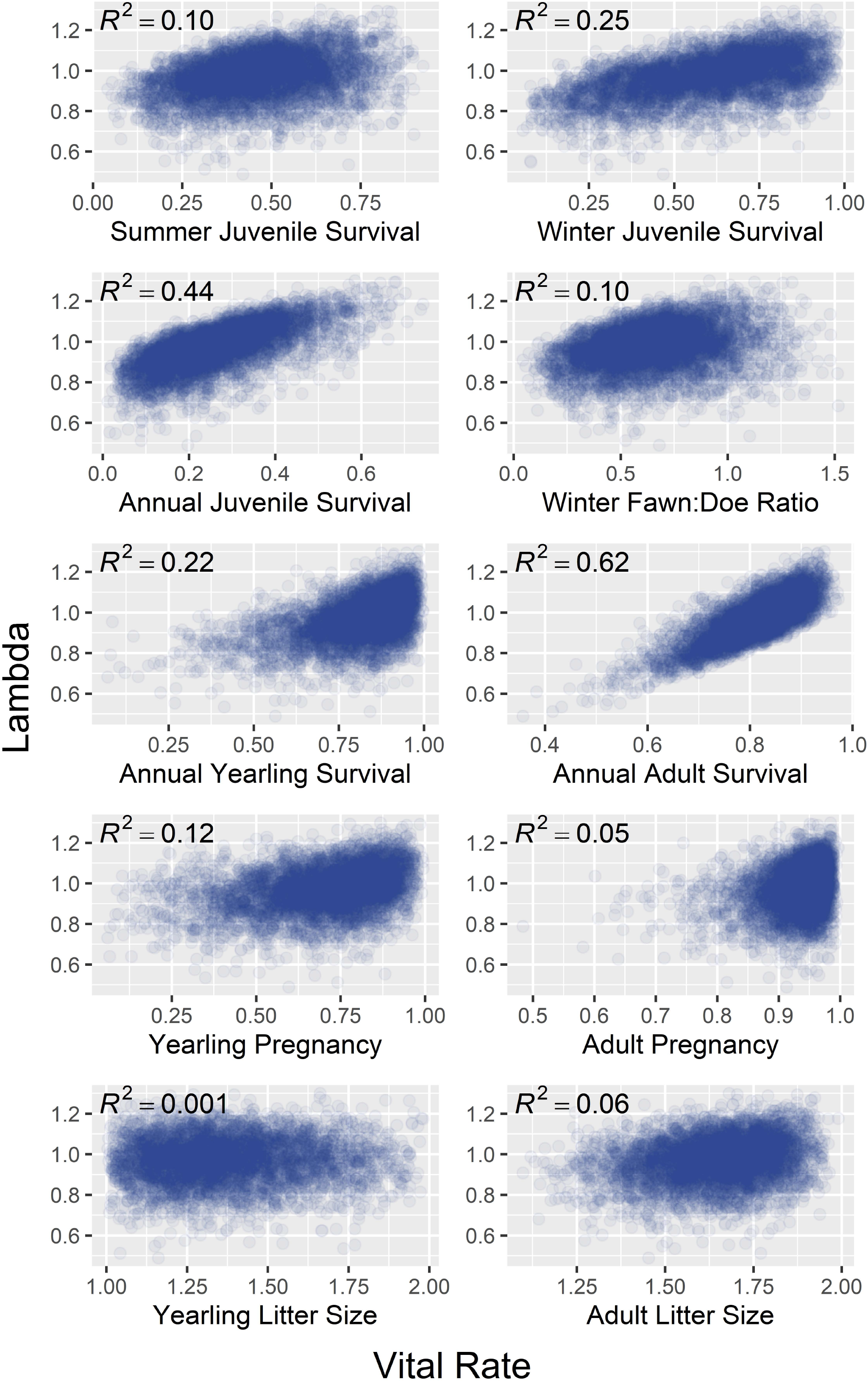
Results from life-stage simulation analysis on mule and black-tailed deer using vital rates summarized across their geographic distribution. Each scatterplot relates population growth (lambda) to an individual vital rate, wherein each datapoint reflects a random draw from a multivariate logit-normal probability distribution characterizing vital rates. Each set of randomly generated vital rates were then used to populate a matrix model used to calculate lambda. Simple linear regression was used to model lambda as a function of each individual vital rate, and *R^2^* values measure the proportion of variation in lambda explained by each vital rate. In each plot, a random sample of 5,000 of the 50,000 simulated data points are plotted.

For a range of plausible values of key vital rates (juvenile survival, adult survival, and fawn:doe ratio), we calculated the proportion of instances in the simulated dataset that resulted in lambda values ≥ 1. For juvenile survival (either summer, winter, or annual) and fawn:doe ratios, the probability of observing a stable lambda began to plateau at high values of the vital rate (Fig. 6). However, this was not the case for adult female survival, in which the probability of observing a stable lambda continued to increase linearly at high values of the vital rate (Fig. 6, Supporting Information S5: Table S2). In contrast to the juvenile survival rates in which a stable lambda could be achieved across a wide range of values, there was almost no probability of observing a stable population when adult female survival was less than 0.7.

**Figure 6.**
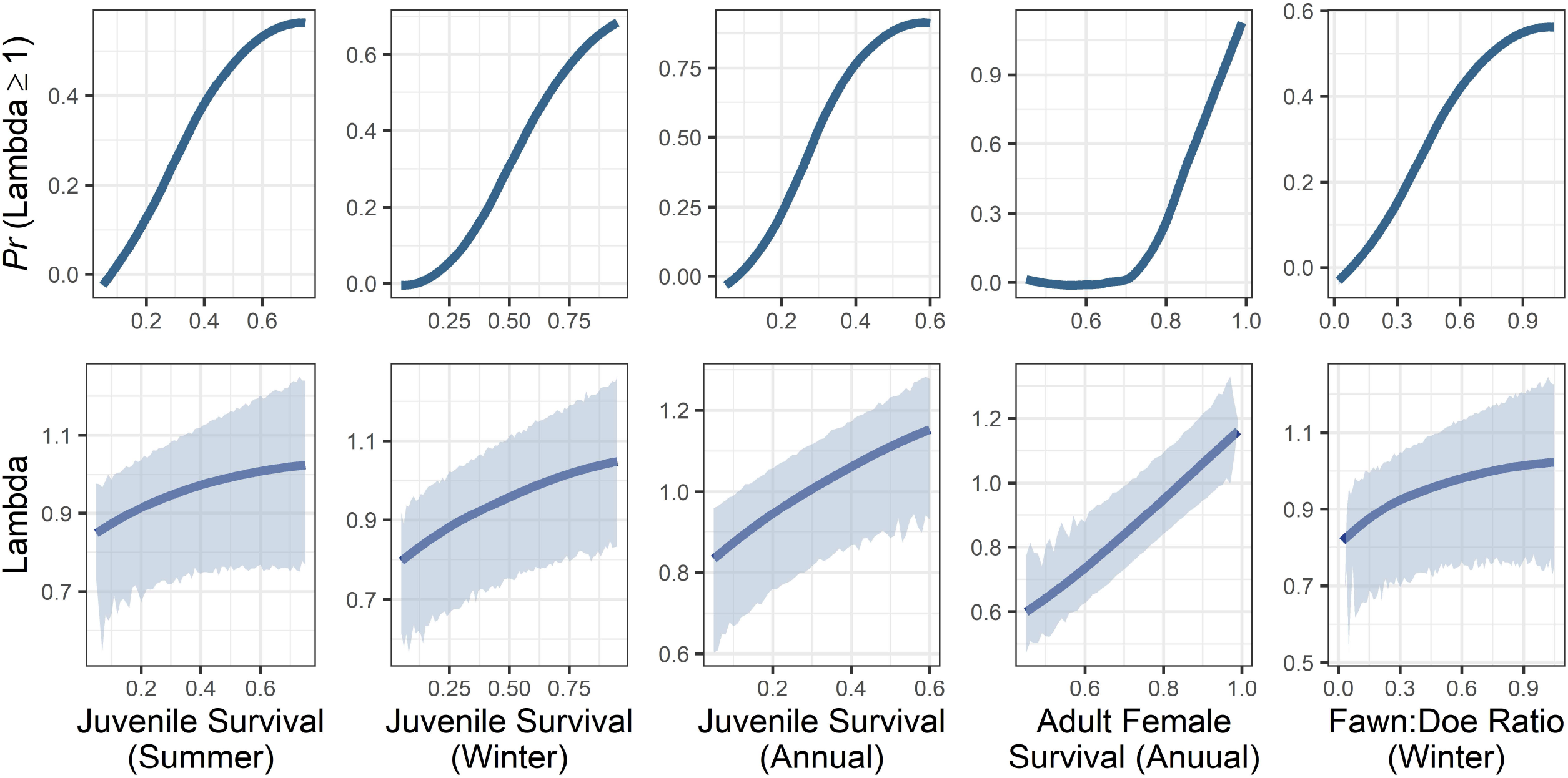
Output from life-stage simulation analysis in mule and black-tailed deer used to make probabilistic statements regarding population growth given key life history parameters. The top row quantifies the probability of observing a stable or increasing lambda given a single vital rate. The bottom row quantifies the expected value of lambda given a single vital rate, wherein the solid line displays the mean prediction of lambda, and the shaded band provides the 95% credible interval for lambda. The winter fawn:doe ratio was constructed from individual vital rates using equation 6 (see text).

The vital rate values that predicted a stable lambda, i.e. the value at which half of the simulations predicted an increasing population and half of the simulations predicted a decreasing population, were 0.48 for summer juvenile survival, 0.62 for winter juvenile survival, and 0.29 for annual juvenile survival, and by extension, values above or below those rates would be indicative of increasing or decreasing populations, respectively. We estimated that adult female survival of 0.84 predicted a stable lambda. Correspondingly, when adult female survival was 0.84, the predicted lambda was 1.0. A start of winter fawn:doe ratio of 71:100 predicted a stable population (Fig. 6).

The mean value of lambda across all 150,000 simulations was 0.975 (*σ* = 0.11), and 61% of these simulations indicated a lambda < 1 (Fig. 7). In our simulations that drew a random value from this distribution of lambda annually for 20 years, at the end of this time period 92% of simulations resulted in a population decline, with a mean decrease of 44%.

**Figure 7:**
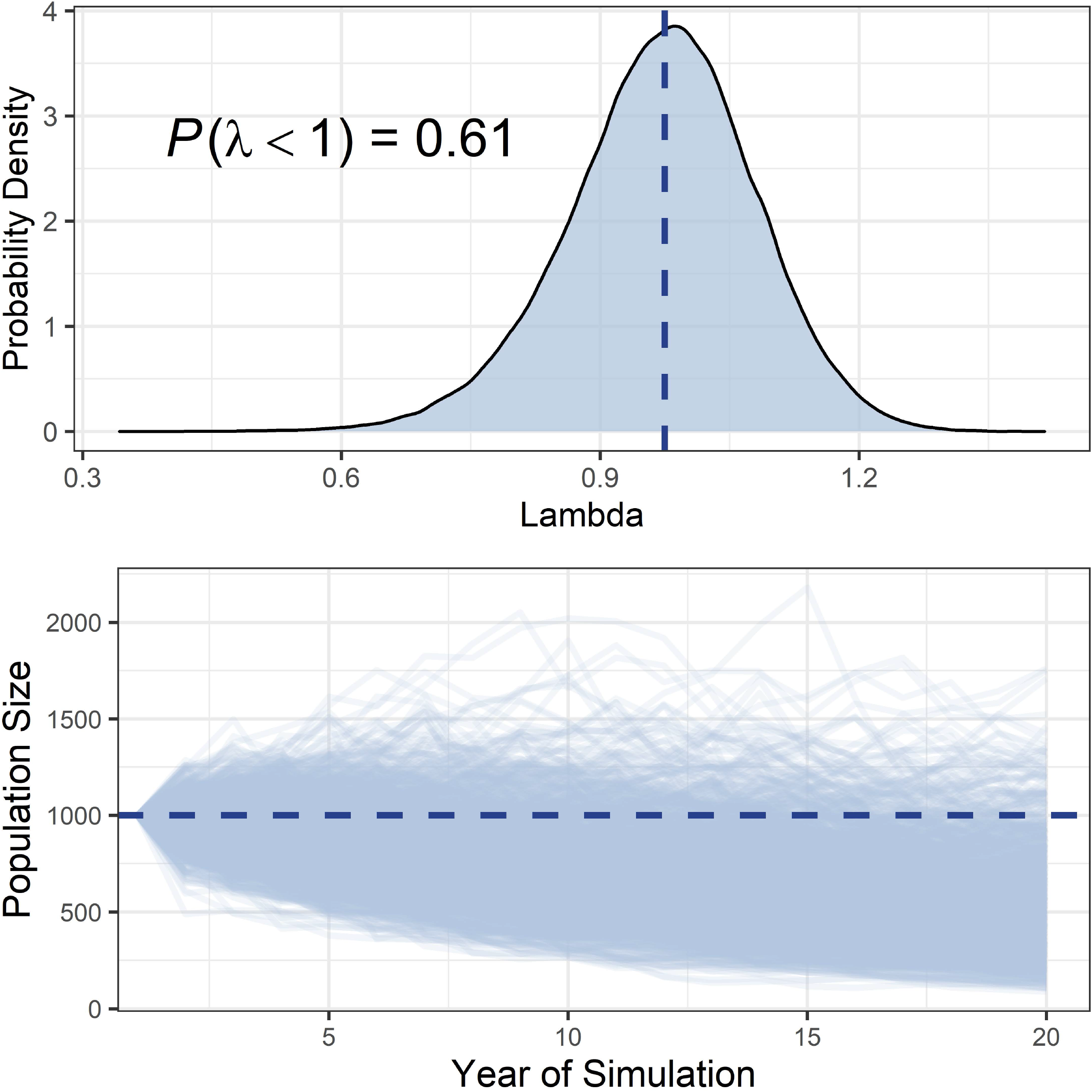
Top panel: the distribution of lambda values as predicted from our matrix model simulations populated by range-wide estimates of mule and black-tailed deer vital rates. The dashed vertical line displays the mean of the distribution (λ = 0.975), and 61% of all simulated lambda values were less than 1. Bottom panel: simulations of 10,000 population trajectories over a period of 20 years, each with a starting population size of 1,000. Each population trajectory was simulated using randomly drawn values of lambda for each year. After 20 years, 92% of populations declined, with an average decrease of 44%.

## DISCUSSION

Our study suggests that mule and black-tailed deer deviate substantially from the paradigm thought to govern the population dynamics of many other large herbivores in which most variation in population growth is driven by the high variability of juvenile survival or recruitment. Instead, we demonstrate that adult female survival, by virtue of being on average lower and more variable than other large ungulates combined with a very high elasticity, determines the majority of interannual variation in lambda. Ungulate species for which both elasticity and LSA results point unequivocally to adult female survival as having the strongest contribution to population growth are rare (but see DeCesare et al. (2012) and Eacker et al. (2017)). While summer and overwinter juvenile survival combined to explain 44% of the variation in lambda, adult survival explained more variation by nearly 20 percentage points. This is in stark contrast to elk wherein a range-wide analysis similar to ours found that three quarters of the variation in lambda was explained by calf survival, and far less by adult female survival (Raithel et al. 2007). Despite mule and black-tailed deer having a much greater reproductive capacity than elk because they can first breed as yearlings and can have larger litter sizes, fecundity contributed little to lambda both in terms of elasticity and its predictive ability.

Our range-wide compilation of mule and black-tailed deer vital rates that was needed to characterize the components of population growth for our simulations generally agreed with the results of the literature review by Forrester and Wittmer (2013). However, one difference was our estimation that adult female survival averaged 0.826 whereas that study estimated a mean of 0.84. While this difference in survival may seem trivial, our simulations suggest that this discrepancy could be the difference between a decreasing and stable population (lambda = 0.98 and 1.0, respectively). Although our meta-analysis used the same sources as Forrester and Wittmer (2013) our review added many additional studies that occurred during the last decade, which is probably the reason for the difference.

Our modeling of the correlations among vital rates showed that most of the significant relationships between vital rates were positive. This pattern has also been observed in other species of ungulates (Coulson et al. 2005, Lukacs et al. 2009) and other taxa (Sæther and Bakke 2000, Fay et al. 2022). Our observation that reproductive parameters positively correlated with one another suggests that in years in which nutritional condition was high, all reproductive parameters would benefit, but this relationship would most likely affect yearling females as annual variation in reproductive parameters, particularly pregnancy rates, is reduced in adults (Monteith et al. 2023). This could be considered a bet hedging strategy, in which during the best years of nutrition, deer would invest heavily into reproductive output and reduce output in poorer years (Sæther and Bakke 2000). We found that adult pregnancy rate was significantly positively correlated with adult survival rate suggesting when nutritional limitation is so great that it reduces pregnancy rate, it could also lead to adult mortality due to malnutrition. The fact that all significant correlations with adult female survival means that focusing management actions on improving adult female survival will likely generate the greatest benefit not only because this has the greatest effect on population growth, but other correlated vital rates (e.g., fawn survival) will likely also improve to further increase population performance.

It was not surprising that overwinter juvenile survival positively correlated with both yearling and adult survival, and we similarly detected correlations between yearling and adult survival (although the latter was not significant). An unexpected result was that summer juvenile survival did not correlate with annual adult survival which we expected because mortality of adult females would logically also result in the death of any neonates they birthed. More surprisingly, however, was the finding that summer and winter juvenile survival were negatively correlated. There are several reasons why this might be the case. First, if all fawns survived the summer, any winter mortality would mean winter survival would necessarily be lower and would lead to a negative correlation. Alternatively, if the smallest fawns all died in summer (Bishop et al. 2009, Hurley et al. 2011, Shallow et al. 2015), the sample surviving until winter should have a larger body mass and be better suited to survive. This conclusion is well supported by the effects of maternal condition on juvenile survival in mule deer (Bishop et al. 2009, Monteith et al. 2014, Lamb et al. 2023, Monteith et al. 2023). This finding also has implications for the effects of summer fawn predation on mule deer population growth. The largest reported cause of fawn mortality in the summer is predation (Forrester and Wittmer 2013). Our finding that low fawn survival in summer is associated with higher winter survival, along with results from predator removal studies that show no increase in mule deer population growth (e.g., Hurley et al. (2011), also see Forrester and Wittmer (2013)), provides further evidence that summer fawn predation may be mostly compensatory. The negative correlation between summer and winter juvenile survival may also be mediated by climate, in which cold conditions during parturition may cause neonates to die from hypothermia (Linnell et al. 1995, Grovenburg et al. 2012), but these same conditions may be associated with longer access to nutritious forage in summer and would support good body condition entering winter (Pettorelli et al. 2007). Nonetheless, in situations where reproduction or fawn survival is reduced, it is likely related to nutritional issues (Bishop et al. 2009, Monteith et al. 2014) and habitat improvement projects will likely provide the greatest benefit to the population.

As part of the life-stage simulation analysis, we populated matrix models 150,000 times using plausible simulated vital rate values which allowed us to predict both lambda and the probability of observing a growing population as a function of individual vital rate values. However, although our hope was that monitoring a single vital rate could be used to forecast population growth, there was substantial uncertainty in lambda estimates from even the vital rates with the greatest predictive ability. For example, when adult female survival was 84%, we predicted lambda would be 1.0, though with a 95% credible interval of 0.88–1.14. A more definitive finding was that an adult female survival rate of 0.7 or lower would cause populations of mule or black-tailed deer to decrease with near certainty, even if other vital rates were high. While survival rates > 0.7 are rare given that adult female survival should be buffered from environmental variation (Gaillard and Yoccoz 2003), it was observed multiple times in our literature review. Our estimation that on average, a winter fawn:doe ratio of 71:100 would be needed to observe a stable population was higher than 66:100 as estimated by Unsworth et al. (1999); however, that study assumed a higher adult survival rate, and additionally, the relationship we observed had wide credible intervals. Despite their ease of collection, the fact that winter fawn:doe ratios only explained 10% of the variation in lambda, and that a stable population could be observed across nearly all plausible values of fawn:doe ratios, suggests their utility in forecasting population growth is limited, but can provide a useful input to integrated population models currently employed by many wildlife management agencies.

Our finding that the range-wide average vital rates for mule and black-tailed deer predict a lambda of 0.975, or a 2.5% decline per year, is not a positive sign for the species. Further, only 8% of our simulations of population trajectories based on plausible lambda values were indicative of an increasing population over a 20-year time period, with an average decrease of 44% during that time. We caution, however, that this assessment could appear overly bleak if the studies reporting vital rates that formed the basis of our analysis were biased low in some way. This could occur if poorly-performing populations are allocated more resources to be studied by state wildlife agencies and this resulted in a reporting bias. However, our results are broadly consistent with observed declining trends in black-tailed and mule deer across their range (Jensen et al. 2023). For example, of 11 states and provinces reporting estimates of black-tailed and mule deer population estimates between 2000 and 2020 (Jensen et al. 2023), only 17% reported population increases over that time, with an estimated regional population decline of 25%. Of course, not all mule and black-tailed deer populations are currently experiencing declines and further studies should investigate the factors associated with stable and increasing populations and report these vital rates in the literature. Our findings indicate that whether or not mule and black-tailed deer populations will increase or decline will in large part be determined by future adult female survival rates.

## MANAGEMENT IMPLICATIONS

Managers often focus on mule deer life stages of reproduction or juvenile survival, but our results indicate the strong need to monitor and manage adult female survival because it explains the majority of variation in population growth. Moreover, even small changes in adult female survival result in disproportionately large changes in population growth, a pattern we quantified for management consideration. In areas with declining mule deer populations, managers should investigate adult female survival rates to determine if they are at a level that is likely to produce a declining population (i.e., survival below 84%) and if so, use cause-specific mortality evidence to guide management actions. Monitoring adult survival can be done through use of satellite collars or similar technologies, in which a large number of adult females can be used as an unbiased, robust estimate of population-level survival. Management can likewise focus on a wide range of factors to improve adult female survival in a holistic manner; these include improvements in nutrition and other habitat factors; reduction in legal and illegal hunter harvest; mitigation of the many diverse sources of vehicle-caused mortality on roads and highways; and accounting for or managing effects of predation in relation to all other factors. In populations where adult female survival is sufficiently high (e.g., >84%), managers should then investigate other vital rates (e.g., fawn survival and reproduction) to determine if they are responsible for reduced population performance. In situations where reproduction or fawn survival is reduced, it is likely related to nutritional issues (Bishop et al. 2009, Monteith et al. 2014) and habitat improvement projects will likely provide the greatest benefit to the population.

Our probability-based estimates of population growth and associated uncertainty provided by credible intervals allow managers to carefully interpret their monitoring estimates of adult female survival or any other vital-rate estimates in relation to population growth, and to evaluate how well management improvements result in desired changes in population growth or trends. Given the range-wide, long-term declines in populations of mule deer, more holistic approaches to monitoring and management of adult survival are needed to inform and implement best management practices for recovery.

## REFERENCES

Bender, L. C., L. A. Lomas, and J. Browning. 2007. Condition, survival, and cause specific mortality of adult female mule deer in north central New Mexico. The Journal of Wildlife Management 71:1118–1124.

Bishop, C. J., G. C. White, D. J. Freddy, B. E. Watkins, and T. R. Stephenson. 2009. Effect of enhanced nutrition on mule deer population rate of change. Wildlife Monographs 172:1–28.

Caswell, H. 1978. A general formula for the sensitivity of population growth rate to changes in life history parameters. Theoretical Population Biology 14:215–230.

Caswell, H. 1989. Analysis of life table response experiments I. Decomposition of effects on population growth rate. Ecological Modelling 46:221–237.

Coulson, T., J.-M. Gaillard, and M. Festa-Bianchet. 2005. Decomposing the variation in population growth into contributions from multiple demographic rates. Journal of Animal Ecology:789–801.

Crowder, L. B., D. T. Crouse, S. S. Heppell, and T. H. Martin. 1994. Predicting the impact of turtle excluder devices on loggerhead sea turtle populations. Ecological Applications 4:437–445.

DeCesare, N. J., M. Hebblewhite, M. Bradley, K. G. Smith, D. Hervieux, and L. Neufeld. 2012. Estimating ungulate recruitment and growth rates using age ratios. The Journal of Wildlife Management 76:144–153.

Eacker, D. R., P. M. Lukacs, K. M. Proffitt, and M. Hebblewhite. 2017. Assessing the importance of demographic parameters for population dynamics using Bayesian integrated population modeling. Ecological Applications 27:1280–1293.

Fay, R., S. Hamel, M. Van de Pol, J. M. Gaillard, N. G. Yoccoz, P. Acker, M. Authier, B. Larue, C. Le Coeur, and K. R. Macdonald. 2022. Temporal correlations among demographic parameters are ubiquitous but highly variable across species. Ecology Letters 25:1640–1654.

Forrester, T. D., and H. U. Wittmer. 2013. A review of the population dynamics of mule deer and black tailed deer Odocoileus hemionus in N orth A merica. Mammal Review 43:292–308.

Gaillard, J.-M., M. Festa-Bianchet, N. Yoccoz, A. Loison, and C. Toigo. 2000. Temporal variation in fitness components and population dynamics of large herbivores. Annual Review of Ecology and Systematics 31:367–393.

Gaillard, J.-M., M. Festa-Bianchet, and N. G. Yoccoz. 1998. Population dynamics of large herbivores: variable recruitment with constant adult survival. Trends in Ecology & Evolution 13:58–63.

Gaillard, J.-M., and N. G. Yoccoz. 2003. Temporal variation in survival of mammals: a case of environmental canalization? Ecology 84:3294–3306.

Gilbert, S. L., K. J. Hundertmark, M. S. Lindberg, D. K. Person, and M. S. Boyce. 2020. The importance of environmental variability and transient population dynamics for a northern ungulate. Frontiers in Ecology and Evolution 8:531027.

Grovenburg, T. W., R. W. Klaver, and J. A. Jenks. 2012. Survival of white tailed deer fawns in the grasslands of the northern Great Plains. The Journal of Wildlife Management 76:944–956.

Hurley, M. A., J. W. Unsworth, P. Zager, M. Hebblewhite, E. O. Garton, D. M. Montgomery, J. R. Skalski, and C. L. Maycock. 2011. Demographic response of mule deer to experimental reduction of coyotes and mountain lions in southeastern Idaho. Wildlife Monographs 178:1–33.

Jensen, W. F., V. C. Bleich, and D. G. Whittaker. 2023. Historical trends in black-tailed deer, mule deer, and their habitats. Pages 25-42 in Ecology and Management of Black-tailed and Mule Deer of North America. CRC Press.

Lamb, S., B. R. McMillan, M. van de Kerk, P. B. Frandsen, K. R. Hersey, and R. Larsen. 2023. From conception to recruitment: Nutritional condition of the dam dictates the likelihood of success in a temperate ungulate. Frontiers in Ecology and Evolution 11:206.

Lefkovitch, L. P. 1965. The study of population growth in organisms grouped by stages. Biometrics:1–18.

Leslie, P. H. 1945. On the use of matrices in certain population mathematics. Biometrika 33:183–212.

Lewis, E. G. 1942. On the Generation and Growth of a Population. Sankhyā: The Indian Journal of Statistics 6:93–96.

Linnell, J. D., R. Aanes, and R. Andersen. 1995. Who killed Bambi? The role of predation in the neonatal mortality of temperate ungulates. Wildlife Biology 1:209–223.

Lukacs, P. M., and J. J. Nowak. 2023. Modeling Population Dynamics of Black-tailed and Mule Deer. Pages 95-102 in Ecology and Management of Black-tailed and Mule Deer of North America. CRC Press.

Lukacs, P. M., G. C. White, B. E. Watkins, R. H. Kahn, B. A. Banulis, D. J. Finley, A. A. Holland, J. A. Martens, and J. Vayhinger. 2009. Separating components of variation in survival of mule deer in Colorado. The Journal of Wildlife Management 73:817–826.

Marescot, L., T. D. Forrester, D. S. Casady, and H. U. Wittmer. 2015. Using multistate capture–mark–recapture models to quantify effects of predation on age-specific survival and population growth in black-tailed deer. Population Ecology 57:185–197.

Miura, S., and K. Tokida. 2009. Management strategy of sika deer based on sensitivity analysis. Pages 453–472 in Sika deer: biology and management of native and introduced populations. Springer.

Monteith, K. L., V. C. Bleich, T. R. Stephenson, B. M. Pierce, M. M. Conner, J. G. Kie, and R. T. Bowyer. 2014. Life history characteristics of mule deer: effects of nutrition in a variable environment. Wildlife Monographs 186:1–62.

Monteith, K. L., T. N. LaSharr, C. J. Bishop, T. R. Stephenson, K. M. Stewart, and L. A. Shipley. 2023. Digestive Physiology and Nutrition. Pages 71-94 in Ecology and Management of Black-tailed and Mule Deer of North America. CRC Press.

Pettorelli, N., F. Pelletier, A. v. Hardenberg, M. Festa-Bianchet, and S. D. Côté. 2007. Early onset of vegetation growth vs. rapid green up: Impacts on juvenile mountain ungulates. Ecology 88:381–390.

Pfister, C. A. 1998. Patterns of variance in stage-structured populations: evolutionary predictions and ecological implications. Proceedings of the National Academy of Sciences 95:213–218.

Raithel, J. D., M. J. Kauffman, and D. H. Pletscher. 2007. Impact of spatial and temporal variation in calf survival on the growth of elk populations. The Journal of Wildlife Management 71:795–803.

Sæther, B.-E., and Ø. Bakke. 2000. Avian life history variation and contribution of demographic traits to the population growth rate. Ecology 81:642–653.

Shallow, J. R., M. A. Hurley, K. L. Monteith, and R. T. Bowyer. 2015. Cascading effects of habitat on maternal condition and life-history characteristics of neonatal mule deer. Journal of Mammalogy 96:194–205.

Unsworth, J. W., D. F. Pac, G. C. White, and R. M. Bartmann. 1999. Mule deer survival in Colorado, Idaho, and Montana. The Journal of Wildlife Management:315–326.

Wisdom, M. J., L. S. Mills, and D. F. Doak. 2000. Life stage simulation analysis: estimating vital rate effects on population growth for conservation. Ecology 81:628–641.

