## Supporting Information S1 for "Variability and correlations among vital rates and their influence on population growth in mule and black-tailed deer"

**SUPPORTING INFORMATION S1: METHODS AND SUMMARY OF LITERATURE REVIEW OF BLACK-TAILED AND MULE DEER VITAL RATES**

*The Journal of Wildlife Management*

**METHODS**

We queried Google Scholar with all combinations of “mule deer”, “black-tailed deer”, or “*Odocoileus hemionus*” and “vital rate”, “demographic rate”, “population dynamics”, “survival”, “reproduction”, “pregnancy”, “litter size”, “fecundity”, and “mortality.” We only used estimates in which a standard error associated with the point estimate was provided or in which we could derive one using the reported sample size or the 95% confidence interval. If a study did not provide a standard error but included a 95% confidence interval (CI), we imputed the standard error using $SE= \frac{(upper CI-lower CI)}{3.92}$. In cases in which the 95% CI was only provided in a plot, we extracted the 95% CI bounds from the plot image using Graph Grabber 2.0.2 (Quintessa 2021; https://www.quintessa.org/software/downloads-and-demos/graph-grabber-2.0.2). If neither the standard error nor the 95% CI was provided but the sample size was reported, we used a beta-binomial model to estimate a standard error using the reported sample size as the number of trials and number of events (e.g. number surviving, pregnant, etc.) as the number of successes.

**RESULTS**

We used 72 data sources reporting a total of 706 annual point estimates across 8 vital rates, plus an additional 16 point estimates for annual juvenile survival (i.e., without separate summer and winter juvenile survival estimates). Of the 72 data sources, 60 were conducted on mule deer, 11 on black-tailed deer, and 1 on hybrid mule and black-tailed in their introgression zone. These studies represented 14 states and 2 Canadian provinces with most studies conducted during the 2000s (Fig. S1). The most commonly-reported vital rates were adult female survival and winter juvenile survival, and the least common were yearling survival and yearling litter size (Fig. S2).

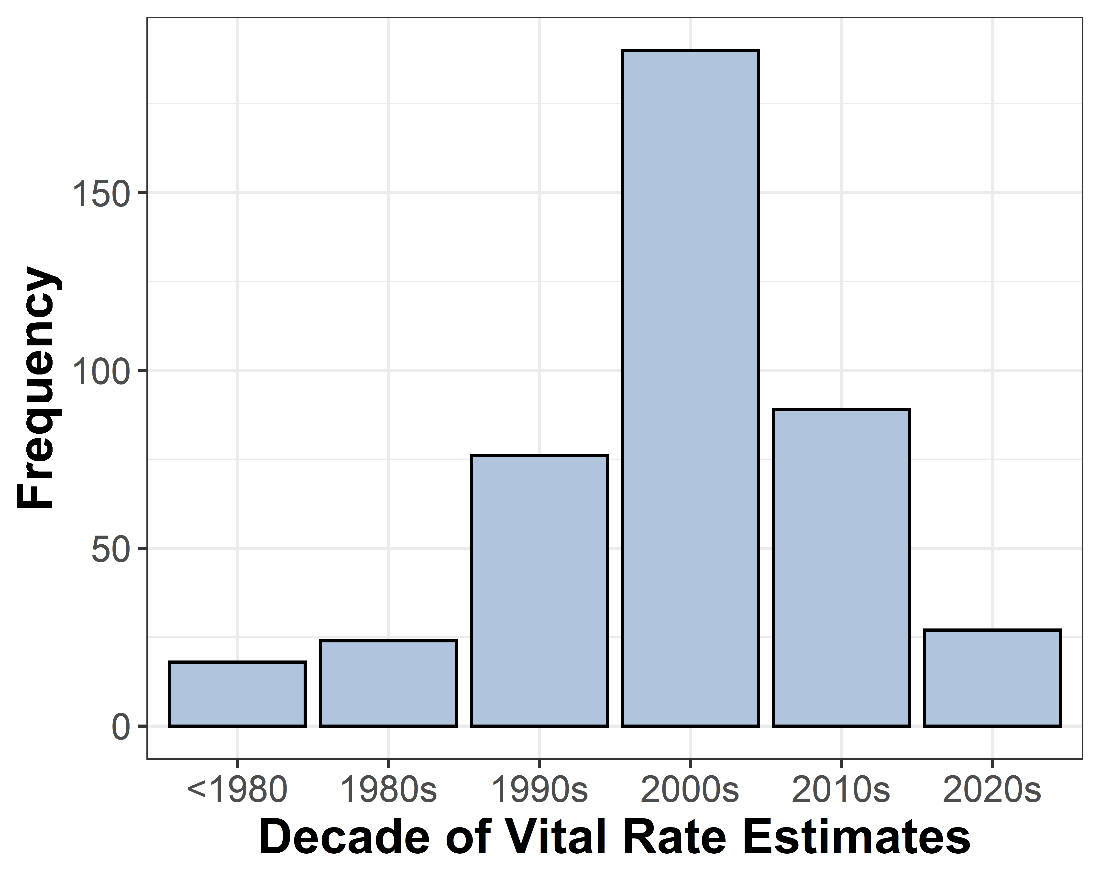

**Fig. S1**: Number of studies reporting vital rates per decade used in our meta-analysis of mule and black-tailed deer vital rates. If the study was conducted over a period that spanned decades, we plotted the most recent decade of the study.

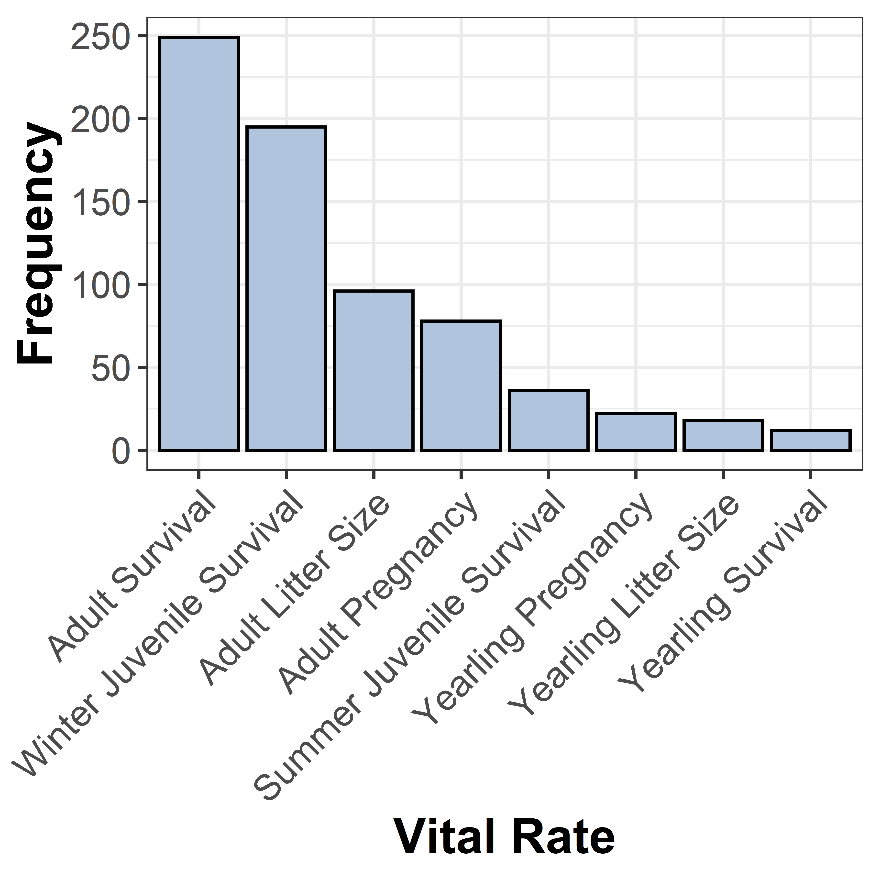

**Fig. S2**: Number of annual point estimates of each vital rate used in our meta-analysis of mule and black-tailed deer.
