## Supporting Information S2 for "Variability and correlations among vital rates and their influence on population growth in mule and black-tailed deer"

**SUPPORTING INFORMATION S2: BLACK-TAILED AND MULE DEER VITAL RATE ESTIMATES USED IN META-ANALYSIS AND THEIR SOURCES**

*The Journal of Wildlife Management*

| Vital Rate | Estimate | SE | Species | State/  Province | Study Area | Year | Study |
| --- | --- | --- | --- | --- | --- | --- | --- |
| Adult female survival (annual) | 0.63 | 0.09 | Mule deer | NM |  | 2002 | Bender et al. 2007/ Lomas and Bender 2007 |
| Adult female survival (annual) | 0.9 | 0.05 | Mule deer | NM |  | 2003 | Bender et al. 2007/ Lomas and Bender 2007 |
| Adult female survival (annual) | 0.91 | 0.04 | Mule deer | NM |  | 2004 | Bender et al. 2007/ Lomas and Bender 2007 |
| Adult female survival (annual) | 0.42 | 0.14 | Mule deer | NM |  | 2006 | Bender et al. 2011 |
| Adult female survival (annual) | 0.78 | 0.08 | Mule deer | NM |  | 2007 | Bender et al. 2011 |
| Adult female survival (annual) | 0.71 | 0.11 | Mule deer | NM |  | 2008 | Bender et al. 2011 |
| Adult female survival (annual) | 0.863 | 0.052 | Mule deer | NM |  | 2004 | Bender et al. 2012 |
| Adult female survival (annual) | 0.829 | 0.051 | Mule deer | NM |  | 2005 | Bender et al. 2012 |
| Adult female survival (annual) | 0.74 | 0.059 | Mule deer | NM |  | 2006 | Bender et al. 2012 |
| Adult female survival (annual) | 0.825 | 0.055 | Mule deer | NM |  | 2007 | Bender et al. 2012 |
| Adult female survival (annual) | 0.804 | 0.058 | Mule deer | NM |  | 2008 | Bender et al. 2012 |
| Adult female survival (annual) | 0.774 | 0.086 | Mule deer | NM |  | 2009 | Bender et al. 2012 |
| Adult female survival (annual) | 0.659 | 0.08 | Mule deer | ID | Bennett | 1993-94 | Bishop et al. 2005 |
| Adult female survival (annual) | 0.884 | 0.055 | Mule deer | ID | Bennett | 1994-95 | Bishop et al. 2005 |
| Adult female survival (annual) | 0.817 | 0.066 | Mule deer | ID | Bennett | 1995-96 | Bishop et al. 2005 |
| Adult female survival (annual) | 0.942 | 0.043 | Mule deer | ID | Bennett | 1996-97 | Bishop et al. 2005 |
| Adult female survival (annual) | 0.853 | 0.077 | Mule deer | ID | Blacks Creek | 1993-94 | Bishop et al. 2005 |
| Adult female survival (annual) | 0.93 | 0.048 | Mule deer | ID | Blacks Creek | 1994-95 | Bishop et al. 2005 |
| Adult female survival (annual) | 0.972 | 0.03 | Mule deer | ID | Blacks Creek | 1995-96 | Bishop et al. 2005 |
| Adult female survival (annual) | 0.839 | 0.062 | Mule deer | ID | Blacks Creek | 1996-97 | Bishop et al. 2005 |
| Adult female survival (annual) | 0.722 | 0.095 | Mule deer | ID | Owyhee | 1993-94 | Bishop et al. 2005 |
| Adult female survival (annual) | 0.765 | 0.103 | Mule deer | ID | Owyhee | 1994-95 | Bishop et al. 2005 |
| Adult female survival (annual) | 0.662 | 0.111 | Mule deer | ID | Owyhee | 1995-96 | Bishop et al. 2005 |
| Adult female survival (annual) | 0.686 | 0.107 | Mule deer | ID | Owyhee | 1996-97 | Bishop et al. 2005 |
| Adult female survival (annual) | 0.867 | 0.023 | Mule deer | CO | Control | 2002-2004 | Bishop et al. 2009 |
| Adult female survival (annual) | 0.898 | 0.019 | Mule deer | CO | Treatment | 2002-2004 | Bishop et al. 2009 |
| Adult female survival (annual) | 0.837 | 0.014 | Mule deer | CA | Casa Diablo | 1986-1994 | Bleich and Taylor 1998 |
| Adult female survival (annual) | 0.884 | 0.014 | Mule deer | CA | East Walker | 1986-1994 | Bleich and Taylor 1998 |
| Adult female survival (annual) | 0.717 | 0.022 | Mule deer | CA | Inyo Mountains | 1986-1994 | Bleich and Taylor 1998 |
| Adult female survival (annual) | 0.777 | 0.018 | Mule deer | CA | Mono Lake | 1986-1994 | Bleich and Taylor 1998 |
| Adult female survival (annual) | 0.643 | 0.01 | Mule deer | CA | West Walker | 1986-1994 | Bleich and Taylor 1998 |
| Adult female survival (annual) | 0.875 | 0.02 | Mule deer | CA |  | 1993 | Bleich et al. 2006 |
| Adult female survival (annual) | 0.929 | 0.01 | Mule deer | CA |  | 1994 | Bleich et al. 2006 |
| Adult female survival (annual) | 0.769 | 0.014 | Mule deer | CA |  | 1995 | Bleich et al. 2006 |
| Adult female survival (annual) | 0.85 | 0.12 | Mule deer | CA |  | 1996 | Bleich et al. 2006 |
| Adult female survival (annual) | 0.85 | 0.022 | Mule deer | CA |  | 2009-2014 | Bush 2015 |
| Adult female survival (annual) | 0.84 | 0.08 | Black-tailed deer | OR |  | 2012 | Clark et al. 2022 |
| Adult female survival (annual) | 0.83 | 0.07 | Black-tailed deer | OR |  | 2013 | Clark et al. 2022 |
| Adult female survival (annual) | 0.71 | 0.07 | Black-tailed deer | OR |  | 2014 | Clark et al. 2022 |
| Adult female survival (annual) | 0.84 | 0.05 | Black-tailed deer | OR |  | 2015 | Clark et al. 2022 |
| Adult female survival (annual) | 0.83 | 0.05 | Black-tailed deer | OR |  | 2016 | Clark et al. 2022 |
| Adult female survival (annual) | 0.79 | 0.051 | Mule deer | MT | Cabinet-Salish | 2017-2019 | DeCesare et al. 2021 |
| Adult female survival (annual) | 0.77 | 0.048 | Mule deer | MT | Rocky Mountain Front | 2017-2019 | DeCesare et al. 2021 |
| Adult female survival (annual) | 0.75 | 0.051 | Mule deer | MT | Whitefish | 2017-2019 | DeCesare et al. 2021 |
| Adult female survival (annual) | 0.67 | 0.08 | Mule deer | WA | Non wolf | 2013-2016 | Dellinger et al. 2018 |
| Adult female survival (annual) | 0.74 | 0.07 | Mule deer | WA | Wolf | 2013-2016 | Dellinger et al. 2018 |
| Adult female survival (annual) | 0.69 | 0.06 | Black-tailed deer | AK |  | 1997-2000 | Farmer et al. 2006 |
| Adult female survival (annual) | 0.71 | 0.07 | Black-tailed deer | CA |  | 2009-2012 | Forrester and Wittmer 2019; Marescot et al. 2015 |
| Adult female survival (annual) | 0.89 | 0.07 | Black-tailed deer | AK |  | 2010 | Gilbert et al. 2020 |
| Adult female survival (annual) | 0.85 | 0.08 | Black-tailed deer | AK |  | 2011 | Gilbert et al. 2020 |
| Adult female survival (annual) | 0.95 | 0.04 | Black-tailed deer | AK |  | 2012 | Gilbert et al. 2020 |
| Adult female survival (annual) | 0.813 | 0.028 | Mule deer | WY |  | 1993-95 | Gogan et al. 2019 |
| Adult female survival (annual) | 0.9 | 0.026 | Black-tailed deer | BC |  | 1970-76 | Hatter and Janz 1994/Hatter and McDermott 2021 |
| Adult female survival (annual) | 0.76 | 0.036 | Black-tailed deer | BC |  | 1977-83 | Hatter and Janz 1994/Hatter and McDermott 2021 |
| Adult female survival (annual) | 0.94 | 0.029 | Black-tailed deer | BC |  | 1984-1990 | Hatter and Janz 1994/Hatter and McDermott 2021 |
| Adult female survival (annual) | 0.71 | 0.012 | Black-tailed deer | BC |  | 1991-2000 | Hatter and Janz 1994/Hatter and McDermott 2021 |
| Adult female survival (annual) | 0.84 | 0.011 | Black-tailed deer | BC |  | 2001-2008 | Hatter and Janz 1994/Hatter and McDermott 2021 |
| Adult female survival (annual) | 0.69 | 0.015 | Black-tailed deer | BC |  | 2009-2015 | Hatter and Janz 1994/Hatter and McDermott 2021 |
| Adult female survival (annual) | 0.91 | 0.053 | Black-tailed deer | BC |  | 2016-2019 | Hatter and Janz 1994/Hatter and McDermott 2021 |
| Adult female survival (annual) | 0.93 | 0.017 | Mule deer | TX |  | 2015-2019 | Heffelfinger et al. 2023 |
| Adult female survival (annual) | 0.84 | 0.046 | Mule deer | WA |  | 2018-2021 | Hellesto et al. 2023 |
| Adult female survival (annual) | 0.879 | 0.015 | Mule deer | ID | Overall | 1998 | Hurley et al. 2011 |
| Adult female survival (annual) | 0.922 | 0.013 | Mule deer | ID | Overall | 1999 | Hurley et al. 2011 |
| Adult female survival (annual) | 0.87 | 0.021 | Mule deer | ID | Reference | 1998 | Hurley et al. 2011 |
| Adult female survival (annual) | 0.844 | 0.026 | Mule deer | ID | Reference | 1999 | Hurley et al. 2011 |
| Adult female survival (annual) | 0.937 | 0.019 | Mule deer | ID | Reference | 2000 | Hurley et al. 2011 |
| Adult female survival (annual) | 0.931 | 0.017 | Mule deer | ID | Reference | 2001 | Hurley et al. 2011 |
| Adult female survival (annual) | 0.768 | 0.03 | Mule deer | ID | Reference | 2002 | Hurley et al. 2011 |
| Adult female survival (annual) | 0.887 | 0.022 | Mule deer | ID | Treatment | 1998 | Hurley et al. 2011 |
| Adult female survival (annual) | 0.98 | 0.018 | Mule deer | ID | Treatment | 1999 | Hurley et al. 2011 |
| Adult female survival (annual) | 0.883 | 0.023 | Mule deer | ID | Treatment | 2000 | Hurley et al. 2011 |
| Adult female survival (annual) | 0.862 | 0.021 | Mule deer | ID | Treatment | 2001 | Hurley et al. 2011 |
| Adult female survival (annual) | 0.706 | 0.034 | Mule deer | ID | Treatment | 2002 | Hurley et al. 2011 |
| Adult female survival (annual) | 0.84 | 0.04 | Mule deer | OR |  | 2014 | Jackson et al. 2021 |
| Adult female survival (annual) | 0.83 | 0.04 | Mule deer | OR |  | 2015 | Jackson et al. 2021 |
| Adult female survival (annual) | 0.8 | 0.04 | Mule deer | OR |  | 2016 | Jackson et al. 2021 |
| Adult female survival (annual) | 0.61 | 0.08 | Mule deer | OR |  | 2017 | Jackson et al. 2021 |
| Adult female survival (annual) | 0.81 | 0.04 | Mule deer | OR |  | 2018 | Jackson et al. 2021 |
| Adult female survival (annual) | 0.73 | 0.09 | Mule deer | KS | North | 2018 | Karish 2022 |
| Adult female survival (annual) | 0.93 | 0.09 | Mule deer | KS | North | 2019 | Karish 2022 |
| Adult female survival (annual) | 0.76 | 0.09 | Mule deer | KS | North | 2020 | Karish 2022 |
| Adult female survival (annual) | 0.87 | 0.08 | Mule deer | KS | South | 2018 | Karish 2022 |
| Adult female survival (annual) | 0.67 | 0.08 | Mule deer | KS | South | 2019 | Karish 2022 |
| Adult female survival (annual) | 0.75 | 0.09 | Mule deer | KS | South | 2020 | Karish 2022 |
| Adult female survival (annual) | 0.826 | 0.114 | Mule deer | MT |  | 2014-2016 | Kolar et al. 2018 |
| Adult female survival (annual) | 0.59 | 0.091 | Mule deer | TX |  | 1990 | Lawrence et al. 2004 |
| Adult female survival (annual) | 0.91 | 0.043 | Mule deer | TX |  | 1991 | Lawrence et al. 2004 |
| Adult female survival (annual) | 0.88 | 0.046 | Mule deer | TX |  | 1992 | Lawrence et al. 2004 |
| Adult female survival (annual) | 0.76 | 0.051 | Mule deer | SD | Black Hills | 2000 | Lindbloom 2021 |
| Adult female survival (annual) | 0.83 | 0.036 | Mule deer | SD | Grand River | 2000 | Lindbloom 2021 |
| Adult female survival (annual) | 0.8 | 0.036 | Mule deer | SD | Upper Missouri | 2000 | Lindbloom 2021 |
| Adult female survival (annual) | 0.84 | 0.036 | Mule deer | SD | White River | 2000 | Lindbloom 2021 |
| Adult female survival (annual) | 0.897 | 0.049 | Mule deer | CO | D-16 | 1999–2000 | Lukacs et al. 2009 |
| Adult female survival (annual) | 0.844 | 0.058 | Mule deer | CO | D-16 | 2000–2001 | Lukacs et al. 2009 |
| Adult female survival (annual) | 0.705 | 0.078 | Mule deer | CO | D-16 | 2001–2002 | Lukacs et al. 2009 |
| Adult female survival (annual) | 0.784 | 0.068 | Mule deer | CO | D-16 | 2002–2003 | Lukacs et al. 2009 |
| Adult female survival (annual) | 0.786 | 0.063 | Mule deer | CO | D-16 | 2003–2004 | Lukacs et al. 2009 |
| Adult female survival (annual) | 0.82 | 0.049 | Mule deer | CO | D-16 | 2004–2005 | Lukacs et al. 2009 |
| Adult female survival (annual) | 0.912 | 0.037 | Mule deer | CO | D-16 | 2005–2006 | Lukacs et al. 2009 |
| Adult female survival (annual) | 0.857 | 0.044 | Mule deer | CO | D-16 | 2006–2007 | Lukacs et al. 2009 |
| Adult female survival (annual) | 0.852 | 0.056 | Mule deer | CO | D-19 | 1997–1998 | Lukacs et al. 2009 |
| Adult female survival (annual) | 0.859 | 0.039 | Mule deer | CO | D-19 | 1998–1999 | Lukacs et al. 2009 |
| Adult female survival (annual) | 0.877 | 0.041 | Mule deer | CO | D-19 | 1999–2000 | Lukacs et al. 2009 |
| Adult female survival (annual) | 0.842 | 0.042 | Mule deer | CO | D-19 | 2000–2001 | Lukacs et al. 2009 |
| Adult female survival (annual) | 0.755 | 0.046 | Mule deer | CO | D-19 | 2001–2002 | Lukacs et al. 2009 |
| Adult female survival (annual) | 0.804 | 0.043 | Mule deer | CO | D-19 | 2002–2003 | Lukacs et al. 2009 |
| Adult female survival (annual) | 0.782 | 0.044 | Mule deer | CO | D-19 | 2003–2004 | Lukacs et al. 2009 |
| Adult female survival (annual) | 0.74 | 0.047 | Mule deer | CO | D-19 | 2004–2005 | Lukacs et al. 2009 |
| Adult female survival (annual) | 0.837 | 0.041 | Mule deer | CO | D-19 | 2005–2006 | Lukacs et al. 2009 |
| Adult female survival (annual) | 0.737 | 0.052 | Mule deer | CO | D-19 | 2006–2007 | Lukacs et al. 2009 |
| Adult female survival (annual) | 0.979 | 0.021 | Mule deer | CO | D-7 | 2001–2002 | Lukacs et al. 2009 |
| Adult female survival (annual) | 0.835 | 0.04 | Mule deer | CO | D-7 | 2002–2003 | Lukacs et al. 2009 |
| Adult female survival (annual) | 0.858 | 0.038 | Mule deer | CO | D-7 | 2003–2004 | Lukacs et al. 2009 |
| Adult female survival (annual) | 0.867 | 0.037 | Mule deer | CO | D-7 | 2004–2005 | Lukacs et al. 2009 |
| Adult female survival (annual) | 0.88 | 0.036 | Mule deer | CO | D-7 | 2005–2006 | Lukacs et al. 2009 |
| Adult female survival (annual) | 0.795 | 0.053 | Mule deer | CO | D-7 | 2006–2007 | Lukacs et al. 2009 |
| Adult female survival (annual) | 0.975 | 0.025 | Mule deer | CO | D-9 | 1998–1999 | Lukacs et al. 2009 |
| Adult female survival (annual) | 0.911 | 0.043 | Mule deer | CO | D-9 | 1999–2000 | Lukacs et al. 2009 |
| Adult female survival (annual) | 0.889 | 0.045 | Mule deer | CO | D-9 | 2000–2001 | Lukacs et al. 2009 |
| Adult female survival (annual) | 0.979 | 0.021 | Mule deer | CO | D-9 | 2001–2002 | Lukacs et al. 2009 |
| Adult female survival (annual) | 0.73 | 0.089 | Mule deer | CO | D-9 | 2002–2003 | Lukacs et al. 2009 |
| Adult female survival (annual) | 0.944 | 0.04 | Mule deer | CO | D-9 | 2003–2004 | Lukacs et al. 2009 |
| Adult female survival (annual) | 0.806 | 0.066 | Mule deer | CO | D-9 | 2004–2005 | Lukacs et al. 2009 |
| Adult female survival (annual) | 0.919 | 0.039 | Mule deer | CO | D-9 | 2005–2006 | Lukacs et al. 2009 |
| Adult female survival (annual) | 0.931 | 0.03 | Mule deer | CO | D-9 | 2006–2007 | Lukacs et al. 2009 |
| Adult female survival (annual) | 0.85 | 0.047 | Mule deer | OR |  | 1992-93 | Mathews and Coggins 1997 |
| Adult female survival (annual) | 0.8 | 0.027 | Black-tailed deer | WA |  | 1989-90 | McCorquodale 1999 |
| Adult female survival (annual) | 0.93 | 0.008 | Black-tailed deer | WA |  | 1990-91 | McCorquodale 1999 |
| Adult female survival (annual) | 0.78 | 0.01 | Black-tailed deer | WA |  | 1991-92 | McCorquodale 1999 |
| Adult female survival (annual) | 0.71 | 0.016 | Black-tailed deer | WA |  | 1992-93 | McCorquodale 1999 |
| Adult female survival (annual) | 0.77 | 0.029 | Black-tailed deer | WA |  | 1993-94 | McCorquodale 1999 |
| Adult female survival (annual) | 0.74 | 0.03 | Black-tailed deer | BC |  | 1982-91 | McNay and Voller 1995 |
| Adult female survival (annual) | 0.87 | 0.025 | Mule deer | CA | Overall | 1997-2009 | Monteith et al. 2014 |
| Adult female survival (annual) | 0.98 | 0.02 | Mule deer | CA |  | 1998 | Monteith et al. 2014 |
| Adult female survival (annual) | 0.946 | 0.032 | Mule deer | CA |  | 1999 | Monteith et al. 2014 |
| Adult female survival (annual) | 0.881 | 0.04 | Mule deer | CA |  | 2000 | Monteith et al. 2014 |
| Adult female survival (annual) | 0.916 | 0.045 | Mule deer | CA |  | 2001 | Monteith et al. 2014 |
| Adult female survival (annual) | 0.839 | 0.043 | Mule deer | CA |  | 2002 | Monteith et al. 2014 |
| Adult female survival (annual) | 0.888 | 0.035 | Mule deer | CA |  | 2003 | Monteith et al. 2014 |
| Adult female survival (annual) | 0.859 | 0.046 | Mule deer | CA |  | 2004 | Monteith et al. 2014 |
| Adult female survival (annual) | 0.78 | 0.065 | Mule deer | CA |  | 2005 | Monteith et al. 2014 |
| Adult female survival (annual) | 0.723 | 0.079 | Mule deer | CA |  | 2006 | Monteith et al. 2014 |
| Adult female survival (annual) | 0.818 | 0.058 | Mule deer | CA |  | 2007 | Monteith et al. 2014 |
| Adult female survival (annual) | 0.98 | 0.024 | Mule deer | CA |  | 2008 | Monteith et al. 2014 |
| Adult female survival (annual) | 0.93 | 0.025 | Mule deer | CA |  | 2005 | Morano 2016 |
| Adult female survival (annual) | 0.93 | 0.025 | Mule deer | CA |  | 2006 | Morano 2016 |
| Adult female survival (annual) | 0.77 | 0.06 | Mule deer | CA |  | 2007 | Morano 2016 |
| Adult female survival (annual) | 0.87 | 0.029 | Mule deer | CA |  | 2008 | Morano 2016 |
| Adult female survival (annual) | 0.788 | 0.023 | Mule deer | OR | Beulah-Malheur | 2016-2022 | ODFW unpublished |
| Adult female survival (annual) | 0.861 | 0.03 | Mule deer | OR | Biggs | 2016-2022 | ODFW unpublished |
| Adult female survival (annual) | 0.787 | 0.011 | Mule deer | OR | Crescent | 2016-2022 | ODFW unpublished |
| Adult female survival (annual) | 0.764 | 0.033 | Mule deer | OR | Fossil-Grizzly | 2016-2022 | ODFW unpublished |
| Adult female survival (annual) | 0.822 | 0.063 | Mule deer | OR | Juniper-Silvies | 2016-2022 | ODFW unpublished |
| Adult female survival (annual) | 0.746 | 0.062 | Mule deer | OR | Keating Creeks | 2016-2022 | ODFW unpublished |
| Adult female survival (annual) | 0.768 | 0.066 | Mule deer | OR | Keno | 2016-2022 | ODFW unpublished |
| Adult female survival (annual) | 0.9 | 0.038 | Mule deer | OR | Klamath Basin | 2016-2022 | ODFW unpublished |
| Adult female survival (annual) | 0.809 | 0.031 | Mule deer | OR | Mid-Columbia | 2016-2022 | ODFW unpublished |
| Adult female survival (annual) | 0.75 | 0.045 | Mule deer | OR | Murderers Creek | 2016-2022 | ODFW unpublished |
| Adult female survival (annual) | 0.775 | 0.062 | Mule deer | OR | North Blues | 2016-2022 | ODFW unpublished |
| Adult female survival (annual) | 0.571 | 0.191 | Mule deer | OR | Northeast | 2016-2022 | ODFW unpublished |
| Adult female survival (annual) | 0.75 | 0.015 | Mule deer | OR | Northside | 2016-2022 | ODFW unpublished |
| Adult female survival (annual) | 0.732 | 0.022 | Mule deer | OR | Ochoco | 2016-2022 | ODFW unpublished |
| Adult female survival (annual) | 0.751 | 0.06 | Mule deer | OR | Southeast | 2016-2022 | ODFW unpublished |
| Adult female survival (annual) | 0.802 | 0.086 | Mule deer | OR | Steens Mnt | 2016-2022 | ODFW unpublished |
| Adult female survival (annual) | 0.698 | 0.072 | Mule deer | OR | Sumpter | 2016-2022 | ODFW unpublished |
| Adult female survival (annual) | 0.785 | 0.075 | Mule deer | OR | Trout Creek | 2016-2022 | ODFW unpublished |
| Adult female survival (annual) | 0.753 | 0.111 | Mule deer | OR | Warner | 2016-2022 | ODFW unpublished |
| Adult female survival (annual) | 0.858 | 0.05 | Mule deer | AZ |  | 2007-2008 | Quintana et al 2016 |
| Adult female survival (annual) | 0.62 | 0.124 | Mule deer | BC |  | 1997 | Robinson et al. 2002 |
| Adult female survival (annual) | 0.68 | 0.12 | Mule deer | BC |  | 1998 | Robinson et al. 2002 |
| Adult female survival (annual) | 0.72 | 0.098 | Mule deer | BC |  | 1999 | Robinson et al. 2002 |
| Adult female survival (annual) | 0.83 | 0.079 | Mule deer | BC |  | 2000 | Robinson et al. 2002 |
| Adult female survival (annual) | 0.815 | 0.01 | Mule deer | OR | Migratory | 2005 | Schuyler et al. 2018 |
| Adult female survival (annual) | 0.817 | 0.01 | Mule deer | OR | Migratory | 2006 | Schuyler et al. 2018 |
| Adult female survival (annual) | 0.817 | 0.01 | Mule deer | OR | Migratory | 2007 | Schuyler et al. 2018 |
| Adult female survival (annual) | 0.816 | 0.01 | Mule deer | OR | Migratory | 2008 | Schuyler et al. 2018 |
| Adult female survival (annual) | 0.817 | 0.01 | Mule deer | OR | Migratory | 2009 | Schuyler et al. 2018 |
| Adult female survival (annual) | 0.821 | 0.01 | Mule deer | OR | Migratory | 2010 | Schuyler et al. 2018 |
| Adult female survival (annual) | 0.817 | 0.01 | Mule deer | OR | Migratory | 2011 | Schuyler et al. 2018 |
| Adult female survival (annual) | 0.755 | 0.026 | Mule deer | OR | Resident | 2005 | Schuyler et al. 2018 |
| Adult female survival (annual) | 0.758 | 0.025 | Mule deer | OR | Resident | 2006 | Schuyler et al. 2018 |
| Adult female survival (annual) | 0.758 | 0.025 | Mule deer | OR | Resident | 2007 | Schuyler et al. 2018 |
| Adult female survival (annual) | 0.757 | 0.025 | Mule deer | OR | Resident | 2008 | Schuyler et al. 2018 |
| Adult female survival (annual) | 0.758 | 0.025 | Mule deer | OR | Resident | 2009 | Schuyler et al. 2018 |
| Adult female survival (annual) | 0.765 | 0.025 | Mule deer | OR | Resident | 2010 | Schuyler et al. 2018 |
| Adult female survival (annual) | 0.759 | 0.025 | Mule deer | OR | Resident | 2011 | Schuyler et al. 2018 |
| Adult female survival (annual) | 0.76 | 0.06 | Mule deer | SA |  | 2006 | Skelton 2010 |
| Adult female survival (annual) | 0.72 | 0.05 | Mule deer | SA |  | 2007 | Skelton 2010 |
| Adult female survival (annual) | 0.86 | 0.08 | Mule deer | SA |  | 2008 | Skelton 2010 |
| Adult female survival (annual) | 0.699 | 0.098 | Mule deer | UT |  | 1979 | Smith 1983 |
| Adult female survival (annual) | 0.866 | 0.058 | Mule deer | UT |  | 1980 | Smith 1983 |
| Adult female survival (annual) | 0.85 | 0.03 | Mule deer | NM |  | 2011 | Sorensen 2015 |
| Adult female survival (annual) | 0.83 | 0.03 | Mule deer | NM |  | 2012 | Sorensen 2015 |
| Adult female survival (annual) | 0.87 | 0.03 | Mule deer | NM |  | 2013 | Sorensen 2015 |
| Adult female survival (annual) | 0.792 | 0.083 | Mule deer | CO |  | 1990 | Symonds 1992 |
| Adult female survival (annual) | 0.857 | 0.059 | Mule deer | CO |  | 1991 | Symonds 1992 |
| Adult female survival (annual) | 0.809 | 0.089 | Mule deer | MT |  | 1977 | Unsworth et al. 1999 |
| Adult female survival (annual) | 0.974 | 0.033 | Mule deer | MT |  | 1978 | Unsworth et al. 1999 |
| Adult female survival (annual) | 0.901 | 0.058 | Mule deer | MT |  | 1979 | Unsworth et al. 1999 |
| Adult female survival (annual) | 0.957 | 0.045 | Mule deer | MT |  | 1980 | Unsworth et al. 1999 |
| Adult female survival (annual) | 0.92 | 0.06 | Mule deer | MT |  | 1981 | Unsworth et al. 1999 |
| Adult female survival (annual) | 0.866 | 0.05 | Mule deer | MT |  | 1982 | Unsworth et al. 1999 |
| Adult female survival (annual) | 0.922 | 0.053 | Mule deer | MT |  | 1983 | Unsworth et al. 1999 |
| Adult female survival (annual) | 0.835 | 0.072 | Mule deer | MT |  | 1984 | Unsworth et al. 1999 |
| Adult female survival (annual) | 0.885 | 0.083 | Mule deer | MT |  | 1985 | Unsworth et al. 1999 |
| Adult female survival (annual) | 0.957 | 0.035 | Mule deer | MT |  | 1986 | Unsworth et al. 1999 |
| Adult female survival (annual) | 0.876 | 0.044 | Mule deer | ID |  | 1992 | Unsworth et al. 1999 |
| Adult female survival (annual) | 0.801 | 0.048 | Mule deer | ID |  | 1993 | Unsworth et al. 1999 |
| Adult female survival (annual) | 0.841 | 0.045 | Mule deer | ID |  | 1994 | Unsworth et al. 1999 |
| Adult female survival (annual) | 0.796 | 0.05 | Mule deer | ID |  | 1995 | Unsworth et al. 1999 |
| Adult female survival (annual) | 0.724 | 0.067 | Mule deer | ID |  | 1996 | Unsworth et al. 1999 |
| Adult female survival (annual) | 0.92 | 0.038 | Mule deer | WA | East Slope Cascades | 2000-2007 | WDFW 2016 |
| Adult female survival (annual) | 0.89 | 0.038 | Mule deer | WA | Northern Rocky Mtns | 2000-2007 | WDFW 2016 |
| Adult female survival (annual) | 0.92 | 0.005 | Mule deer | WA | Columbia Plateau | 2000-2008 | WDFW 2021 |
| Adult female survival (annual) | 0.89 | 0.01 | Mule deer | WA | Okanagan Highlands | 2000-2007 | WDFW 2021 |
| Adult female survival (annual) | 0.848 | 0.071 | Mule deer | CO |  | 1982 | White and Bartmann 1998 |
| Adult female survival (annual) | 0.856 | 0.067 | Mule deer | CO |  | 1983 | White and Bartmann 1998 |
| Adult female survival (annual) | 0.95 | 0.04 | Mule deer | CO |  | 1984 | White and Bartmann 1998 |
| Adult female survival (annual) | 0.911 | 0.042 | Mule deer | CO |  | 1985 | White and Bartmann 1998 |
| Adult female survival (annual) | 0.76 | 0.067 | Mule deer | CO |  | 1986 | White and Bartmann 1998 |
| Adult female survival (annual) | 0.875 | 0.083 | Mule deer | CO |  | 1987 | White and Bartmann 1998 |
| Adult female survival (annual) | 0.833 | 0.108 | Mule deer | CO |  | 1988 | White and Bartmann 1998 |
| Adult female survival (annual) | 0.905 | 0.04 | Mule deer | CO |  | 1989 | White and Bartmann 1998 |
| Adult female survival (annual) | 0.942 | 0.033 | Mule deer | CO |  | 1990 | White and Bartmann 1998 |
| Adult female survival (annual) | 0.763 | 0.05 | Mule deer | CO |  | 1991 | White and Bartmann 1998 |
| Adult female survival (annual) | 0.716 | 0.046 | Mule deer | CO |  | 1992 | White and Bartmann 1998 |
| Adult female survival (annual) | 0.84 | 0.038 | Mule deer | CO |  | 1993 | White and Bartmann 1998 |
| Adult female survival (annual) | 0.878 | 0.035 | Mule deer | CO |  | 1994 | White and Bartmann 1998 |
| Adult female survival (annual) | 0.932 | 0.029 | Mule deer | CO |  | 1995 | White and Bartmann 1998 |
| Adult female survival (annual) | 0.838 | 0.068 | hybrid | CA | Overall | 2017-2020 | Wittmer et al. 2021 |
| Adult female survival (annual) | 0.79 | 0.041 | hybrid | CA |  | 2015-2016 | Wittmer et al. 2021 |
| Adult female survival (annual) | 0.86 | 0.036 | hybrid | CA |  | 2016-2017 | Wittmer et al. 2021 |
| Adult female survival (annual) | 0.84 | 0.036 | hybrid | CA |  | 2017-2018 | Wittmer et al. 2021 |
| Adult female survival (annual) | 0.9 | 0.02 | hybrid | CA |  | 2018-2019 | Wittmer et al. 2021 |
| Adult female survival (annual) | 0.81 | 0.036 | hybrid | CA |  | 2019-2020 | Wittmer et al. 2021 |
| Adult female survival (annual) | 0.77 | 0.028 | Mule deer | BC |  | 2014-2018 | Wright et al. 2020 |
| Adult female survival (annual) | 0.85 | 0.052 | Mule deer | ID | DAU 1 | 2005-2006 | Zager et al. 2007 |
| Adult female survival (annual) | 0.84 | 0.059 | Mule deer | ID | DAU 1 | 2006-2007 | Zager et al. 2007 |
| Adult female survival (annual) | 0.84 | 0.054 | Mule deer | ID | DAU 10 | 2006-2007 | Zager et al. 2007 |
| Adult female survival (annual) | 0.87 | 0.088 | Mule deer | ID | DAU 12 | 2005-2006 | Zager et al. 2007 |
| Adult female survival (annual) | 0.95 | 0.047 | Mule deer | ID | DAU 12 | 2006-2007 | Zager et al. 2007 |
| Adult female survival (annual) | 0.83 | 0.108 | Mule deer | ID | DAU 13 | 2005-2006 | Zager et al. 2007 |
| Adult female survival (annual) | 0.96 | 0.041 | Mule deer | ID | DAU 13 | 2006-2007 | Zager et al. 2007 |
| Adult female survival (annual) | 0.8 | 0.037 | Mule deer | ID | DAU 4 | 2005-2006 | Zager et al. 2007 |
| Adult female survival (annual) | 0.86 | 0.034 | Mule deer | ID | DAU 4 | 2006-2007 | Zager et al. 2007 |
| Adult female survival (annual) | 0.85 | 0.071 | Mule deer | ID | DAU 5 | 2005-2006 | Zager et al. 2007 |
| Adult female survival (annual) | 0.95 | 0.044 | Mule deer | ID | DAU 5 | 2006-2007 | Zager et al. 2007 |
| Adult female survival (annual) | 0.83 | 0.07 | Mule deer | ID | DAU 6 | 2005-2006 | Zager et al. 2007 |
| Adult female survival (annual) | 0.67 | 0.086 | Mule deer | ID | DAU 6 | 2006-2007 | Zager et al. 2007 |
| Adult litter size | 1.49 | 0.06 | Mule deer | NM |  | 2003-2008 | Bender and Hoenes 2018 |
| Adult litter size | 1.32 | 0.11 | Mule deer | NM |  | 2002 | Bender et al. 2007/ Lomas and Bender 2007 |
| Adult litter size | 1.24 | 0.07 | Mule deer | NM |  | 2003 | Bender et al. 2007/ Lomas and Bender 2007 |
| Adult litter size | 1.94 | 0.1 | Mule deer | NM |  | 2004 | Bender et al. 2007/ Lomas and Bender 2007 |
| Adult litter size | 1.394 | 0.24 | Mule deer | CO | Control | 2001-2002 | Bishop et al. 2009 |
| Adult litter size | 1.702 | 0.13 | Mule deer | CO | Control | 2002-2003 | Bishop et al. 2009 |
| Adult litter size | 1.501 | 0.18 | Mule deer | CO | Control | 2003-2004 | Bishop et al. 2009 |
| Adult litter size | 1.534 | 0.2 | Mule deer | CO | Treatment | 2001-2002 | Bishop et al. 2009 |
| Adult litter size | 1.758 | 0.087 | Mule deer | CO | Treatment | 2002-2003 | Bishop et al. 2009 |
| Adult litter size | 1.976 | 0.08 | Mule deer | CO | Treatment | 2003-2004 | Bishop et al. 2009 |
| Adult litter size | 1.56 | 0.178 | Mule deer | CA |  | 1993 | Bleich et al. 2006 |
| Adult litter size | 1.47 | 0.184 | Mule deer | CA |  | 1994 | Bleich et al. 2006 |
| Adult litter size | 1.7 | 0.165 | Mule deer | CA |  | 1995 | Bleich et al. 2006 |
| Adult litter size | 1.82 | 0.304 | Mule deer | CA |  | 1996 | Bleich et al. 2006 |
| Adult litter size | 1.61 | 0.054 | Mule deer | CA |  | 2009-2014 | Bush 2015 |
| Adult litter size | 1.9 | 0.08 | Black-tailed deer | CA |  | 2009-2012 | Forrester and Wittmer 2019; Marescot et al. 2015 |
| Adult litter size | 1.67 | 0.43 | Black-tailed deer | AK |  | 2010 | Gilbert et al. 2020 |
| Adult litter size | 1.36 | 0.35 | Black-tailed deer | AK |  | 2011 | Gilbert et al. 2020 |
| Adult litter size | 1.33 | 0.33 | Black-tailed deer | AK |  | 2012 | Gilbert et al. 2020 |
| Adult litter size | 1.5 | 0.13 | Mule deer | MT |  | 1976 | Hamlin et al. 1984 |
| Adult litter size | 1.52 | 0.096 | Mule deer | MT |  | 1977 | Hamlin et al. 1984 |
| Adult litter size | 1.65 | 0.094 | Mule deer | MT |  | 1978 | Hamlin et al. 1984 |
| Adult litter size | 1.76 | 0.07 | Mule deer | MT |  | 1979 | Hamlin et al. 1984 |
| Adult litter size | 1.68 | 0.099 | Mule deer | MT |  | 1980 | Hamlin et al. 1984 |
| Adult litter size | 1.62 | 0.095 | Mule deer | MT |  | 1981 | Hamlin et al. 1984 |
| Adult litter size | 1.67 | 0.12 | Black-tailed deer | BC | Adam | 1980-81 | Hatter 1988 |
| Adult litter size | 1.63 | 0.09 | Black-tailed deer | BC | Nimpkish | 1980-81 | Hatter 1988 |
| Adult litter size | 1.67 | 0.068 | Black-tailed deer | BC | Northwest | 1980-81 | Hatter 1988 |
| Adult litter size | 1.33 | 0.05 | Mule deer | CA |  | 2013-2016 | Heffelfinger et al. 2018 |
| Adult litter size | 1.62 | 0.097 | Mule deer | ID | Reference | 1998 | Hurley et al. 2011 |
| Adult litter size | 1.77 | 0.094 | Mule deer | ID | Reference | 1999 | Hurley et al. 2011 |
| Adult litter size | 1.84 | 0.069 | Mule deer | ID | Reference | 2000 | Hurley et al. 2011 |
| Adult litter size | 1.81 | 0.099 | Mule deer | ID | Reference | 2001 | Hurley et al. 2011 |
| Adult litter size | 1.68 | 0.107 | Mule deer | ID | Reference | 2002 | Hurley et al. 2011 |
| Adult litter size | 1.62 | 0.097 | Mule deer | ID | Treatment | 1998 | Hurley et al. 2011 |
| Adult litter size | 1.81 | 0.117 | Mule deer | ID | Treatment | 1999 | Hurley et al. 2011 |
| Adult litter size | 1.7 | 0.094 | Mule deer | ID | Treatment | 2000 | Hurley et al. 2011 |
| Adult litter size | 1.7 | 0.102 | Mule deer | ID | Treatment | 2001 | Hurley et al. 2011 |
| Adult litter size | 1.36 | 0.084 | Mule deer | ID | Treatment | 2002 | Hurley et al. 2011 |
| Adult litter size | 1.33 | 0.079 | Mule deer | WA |  | 2003 | Johnstone-Yellin et al 2009 |
| Adult litter size | 1.42 | 0.11 | Mule deer | UT | Antimony | 1954-56 | Julander et al. 1961 |
| Adult litter size | 1.96 | 0.04 | Mule deer | ID | Sublett | 1954-56 | Julander et al. 1961 |
| Adult litter size | 1.66 | 0.06 | Mule deer | UT |  | 2018-2020 | Lamb et al. 2023 |
| Adult litter size | 1.76 | 0.06 | Black-tailed deer | WA |  | 2006 | McCoy and Murphie 2011/McCoy et al. 2014 |
| Adult litter size | 1.68 | 0.066 | Black-tailed deer | WA |  | 2007 | McCoy and Murphie 2011/McCoy et al. 2014 |
| Adult litter size | 1.72 | 0.055 | Black-tailed deer | WA |  | 2008 | McCoy and Murphie 2011/McCoy et al. 2014 |
| Adult litter size | 1.31 | 0.059 | Black-tailed deer | WA |  | 2009 | McCoy and Murphie 2011/McCoy et al. 2014 |
| Adult litter size | 1.355 | 0.104 | Mule deer | CA |  | 1991 | Monteith et al. 2014 |
| Adult litter size | 1.301 | 0.163 | Mule deer | CA |  | 1992 | Monteith et al. 2014 |
| Adult litter size | 1.559 | 0.167 | Mule deer | CA |  | 1993 | Monteith et al. 2014 |
| Adult litter size | 1.472 | 0.125 | Mule deer | CA |  | 1994 | Monteith et al. 2014 |
| Adult litter size | 1.652 | 0.15 | Mule deer | CA |  | 1995 | Monteith et al. 2014 |
| Adult litter size | 1.82 | 0.121 | Mule deer | CA |  | 1996 | Monteith et al. 2014 |
| Adult litter size | 1.617 | 0.141 | Mule deer | CA |  | 1997 | Monteith et al. 2014 |
| Adult litter size | 1.811 | 0.072 | Mule deer | CA |  | 1998 | Monteith et al. 2014 |
| Adult litter size | 1.913 | 0.054 | Mule deer | CA |  | 1999 | Monteith et al. 2014 |
| Adult litter size | 1.754 | 0.054 | Mule deer | CA |  | 2000 | Monteith et al. 2014 |
| Adult litter size | 1.57 | 0.065 | Mule deer | CA |  | 2001 | Monteith et al. 2014 |
| Adult litter size | 1.653 | 0.055 | Mule deer | CA |  | 2002 | Monteith et al. 2014 |
| Adult litter size | 1.687 | 0.049 | Mule deer | CA |  | 2003 | Monteith et al. 2014 |
| Adult litter size | 1.65 | 0.119 | Mule deer | CA |  | 2004 | Monteith et al. 2014 |
| Adult litter size | 1.709 | 0.058 | Mule deer | CA |  | 2005 | Monteith et al. 2014 |
| Adult litter size | 1.727 | 0.07 | Mule deer | CA |  | 2006 | Monteith et al. 2014 |
| Adult litter size | 1.741 | 0.063 | Mule deer | CA |  | 2007 | Monteith et al. 2014 |
| Adult litter size | 1.574 | 0.062 | Mule deer | CA |  | 2008 | Monteith et al. 2014 |
| Adult litter size | 1.594 | 0.064 | Mule deer | CA |  | 2009 | Monteith et al. 2014 |
| Adult litter size | 1.57 | 0.136 | Mule deer | CA |  | 2006 | Morano 2016 |
| Adult litter size | 1.06 | 0.11 | Mule deer | CA |  | 2007 | Morano 2016 |
| Adult litter size | 1.74 | 0.053 | Mule deer | MT |  | 1953-63 | Nellis 1968 and Nellis 1964 |
| Adult litter size | 1.57 | 0.112 | Mule deer | WY | High development | 2012 | Northrup et al. 2021 |
| Adult litter size | 1.68 | 0.119 | Mule deer | WY | High development | 2013 | Northrup et al. 2021 |
| Adult litter size | 1.7 | 0.108 | Mule deer | WY | High development | 2014 | Northrup et al. 2021 |
| Adult litter size | 1.78 | 0.077 | Mule deer | WY | High development | 2015 | Northrup et al. 2021 |
| Adult litter size | 1.87 | 0.115 | Mule deer | WY | Low development | 2012 | Northrup et al. 2021 |
| Adult litter size | 1.83 | 0.113 | Mule deer | WY | Low development | 2013 | Northrup et al. 2021 |
| Adult litter size | 1.67 | 0.124 | Mule deer | WY | Low development | 2014 | Northrup et al. 2021 |
| Adult litter size | 1.71 | 0.104 | Mule deer | WY | Low development | 2015 | Northrup et al. 2021 |
| Adult litter size | 1.97 | 0.029 | Mule deer | SA |  | 2010 | Perera 2012 |
| Adult litter size | 2.06 | 0.018 | Mule deer | SA |  | 2011 | Perera 2012 |
| Adult litter size | 1.7 | 0.07 | Mule deer | CO |  | 1999 | Pojar and Bowden 2004/ Andelt et al. 2004 |
| Adult litter size | 1.77 | 0.067 | Mule deer | AZ |  | 2007-2008 | Quintana et al 2016 |
| Adult litter size | 1.67 | 0.035 | Mule deer | UT |  | 1948-49 | Robinette and Gashwiler 1950 |
| Adult litter size | 1.81 | 0.019 | Mule deer | UT |  | 1950-53 | Robinette et al. 1955 |
| Adult litter size | 1.78 | 0.22 | Mule deer | BC |  | 1997-2000 | Robinson et al. 2002 |
| Adult litter size | 1.96 | 0.04 | Mule deer | ID | Caribou Mountains | 2010-2011 | Shallow et al. 2015 |
| Adult litter size | 1.52 | 0.09 | Mule deer | ID | Salmon River Mountains | 2010-2011 | Shallow et al. 2015 |
| Adult litter size | 1.96 | 0.029 | Mule deer | UT |  | 1979 | Smith 1983 |
| Adult litter size | 1.73 | 0.063 | Mule deer | UT |  | 1980 | Smith 1983 |
| Adult litter size | 1.88 | 0.05 | Mule deer | NM |  | 2011 | Sorensen 2015 |
| Adult litter size | 1.74 | 0.064 | Black-tailed deer | BC |  | 1963-67 | Thomas 1983 |
| Adult litter size | 1.6 | 0.1 | Mule deer | UT |  | 2019-2022 | Turnley et al. 2022 |
| Adult litter size | 1.66 | 0.27 | Mule deer | WA | East Slope Cascades | 2000-2007 | WDFW 2016 |
| Adult litter size | 1.8 | 0.32 | Mule deer | WA | Northern Rocky Mtns | 2000-2007 | WDFW 2016 |
| Adult litter size | 1.44 | 0.122 | Mule deer | WA | Columbia Plateau | 2000-2008 | WDFW 2021 |
| Adult litter size | 1.44 | 0.209 | Mule deer | WA | Okanagan Highlands | 2000-2007 | WDFW 2021 |
| Adult litter size | 1.863 | 0.1 | hybrid | CA | Overall | 2017-2020 | Wittmer et al. 2021 |
| Adult pregnancy | 0.92 | 0.025 | Mule deer | NM |  | 2003-2008 | Bender and Hoenes 2018 |
| Adult pregnancy | 0.67 | 0.07 | Mule deer | NM |  | 2002 | Bender et al. 2007/ Lomas and Bender 2007 |
| Adult pregnancy | 0.935 | 0.019 | Mule deer | CO | Overall | 2002-2004 | Bishop et al. 2009 |
| Adult pregnancy | 0.96 | 0.019 | Mule deer | CA |  | 2009-2014 | Bush 2015 |
| Adult pregnancy | 0.96 | 0.038 | Mule deer | MT | Cabinet-Salish | 2017-2019 | DeCesare et al. 2021 |
| Adult pregnancy | 0.98 | 0.02 | Mule deer | MT | Rocky Mountain Front | 2017-2019 | DeCesare et al. 2021 |
| Adult pregnancy | 0.97 | 0.03 | Mule deer | MT | Whitefish | 2017-2019 | DeCesare et al. 2021 |
| Adult pregnancy | 0.87 | 0.05 | Black-tailed deer | CA |  | 2009-2012 | Forrester and Wittmer 2019; Marescot et al. 2015 |
| Adult pregnancy | 0.986 | 0.014 | Mule deer | UT |  | 2012 | Freeman et al. 2014 |
| Adult pregnancy | 0.966 | 0.016 | Mule deer | CO |  | 2012 | Freeman et al. 2014 |
| Adult pregnancy | 0.95 | 0.043 | Black-tailed deer | AK |  | 2010 | Gilbert et al. 2020 |
| Adult pregnancy | 0.95 | 0.05 | Black-tailed deer | AK |  | 2011 | Gilbert et al. 2020 |
| Adult pregnancy | 0.77 | 0.09 | Black-tailed deer | AK |  | 2012 | Gilbert et al. 2020 |
| Adult pregnancy | 0.95 | 0.013 | Mule deer | UT |  | 2012-2015 | Hall 2018 |
| Adult pregnancy | 0.97 | 0.015 | Mule deer | CA |  | 2013-2016 | Heffelfinger et al. 2018 |
| Adult pregnancy | 0.98 | 0.014 | Mule deer | ID | Overall | 1998 | Hurley et al. 2011 |
| Adult pregnancy | 0.91 | 0.038 | Mule deer | ID | Overall | 1999 | Hurley et al. 2011 |
| Adult pregnancy | 0.94 | 0.06 | Mule deer | KS | North | 2018 | Karish 2022 |
| Adult pregnancy | 0.89 | 0.07 | Mule deer | KS | North | 2019 | Karish 2022 |
| Adult pregnancy | 0.94 | 0.06 | Mule deer | KS | North | 2020 | Karish 2022 |
| Adult pregnancy | 0.94 | 0.06 | Mule deer | KS | South | 2018 | Karish 2022 |
| Adult pregnancy | 0.83 | 0.09 | Mule deer | KS | South | 2019 | Karish 2022 |
| Adult pregnancy | 0.94 | 0.06 | Mule deer | KS | South | 2020 | Karish 2022 |
| Adult pregnancy | 0.95 | 0.02 | Mule deer | UT |  | 2018-2020 | Lamb et al. 2023 |
| Adult pregnancy | 0.87 | 0.06 | Mule deer | TX |  | 1990 | Lawrence et al. 2004 |
| Adult pregnancy | 0.96 | 0.04 | Mule deer | TX |  | 1991 | Lawrence et al. 2004 |
| Adult pregnancy | 0.975 | 0.029 | Mule deer | CA |  | 1991 | Monteith et al. 2014 |
| Adult pregnancy | 0.855 | 0.082 | Mule deer | CA |  | 1992 | Monteith et al. 2014 |
| Adult pregnancy | 0.893 | 0.075 | Mule deer | CA |  | 1993 | Monteith et al. 2014 |
| Adult pregnancy | 0.95 | 0.05 | Mule deer | CA |  | 1994 | Monteith et al. 2014 |
| Adult pregnancy | 0.954 | 0.05 | Mule deer | CA |  | 1995 | Monteith et al. 2014 |
| Adult pregnancy | 0.92 | 0.07 | Mule deer | CA |  | 1996 | Monteith et al. 2014 |
| Adult pregnancy | 0.93 | 0.06 | Mule deer | CA |  | 1997 | Monteith et al. 2014 |
| Adult pregnancy | 0.97 | 0.029 | Mule deer | CA |  | 1998 | Monteith et al. 2014 |
| Adult pregnancy | 0.98 | 0.02 | Mule deer | CA |  | 1999 | Monteith et al. 2014 |
| Adult pregnancy | 0.98 | 0.015 | Mule deer | CA |  | 2000 | Monteith et al. 2014 |
| Adult pregnancy | 0.936 | 0.026 | Mule deer | CA |  | 2001 | Monteith et al. 2014 |
| Adult pregnancy | 0.98 | 0.016 | Mule deer | CA |  | 2002 | Monteith et al. 2014 |
| Adult pregnancy | 0.993 | 0.01 | Mule deer | CA |  | 2003 | Monteith et al. 2014 |
| Adult pregnancy | 0.95 | 0.05 | Mule deer | CA |  | 2004 | Monteith et al. 2014 |
| Adult pregnancy | 0.976 | 0.017 | Mule deer | CA |  | 2005 | Monteith et al. 2014 |
| Adult pregnancy | 0.947 | 0.027 | Mule deer | CA |  | 2006 | Monteith et al. 2014 |
| Adult pregnancy | 0.987 | 0.016 | Mule deer | CA |  | 2007 | Monteith et al. 2014 |
| Adult pregnancy | 0.989 | 0.014 | Mule deer | CA |  | 2008 | Monteith et al. 2014 |
| Adult pregnancy | 0.98 | 0.016 | Mule deer | CA |  | 2009 | Monteith et al. 2014 |
| Adult pregnancy | 0.929 | 0.028 | Mule deer | MT |  | 1953-63 | Nellis 1968 and Nellis 1964 |
| Adult pregnancy | 0.94 | 0.06 | Mule deer | WY | High development | 2009 | Northrup et al. 2021 |
| Adult pregnancy | 0.98 | 0.02 | Mule deer | WY | High development | 2010 | Northrup et al. 2021 |
| Adult pregnancy | 0.95 | 0.041 | Mule deer | WY | High development | 2011 | Northrup et al. 2021 |
| Adult pregnancy | 0.96 | 0.037 | Mule deer | WY | High development | 2012 | Northrup et al. 2021 |
| Adult pregnancy | 0.9 | 0.055 | Mule deer | WY | High development | 2013 | Northrup et al. 2021 |
| Adult pregnancy | 0.93 | 0.045 | Mule deer | WY | High development | 2014 | Northrup et al. 2021 |
| Adult pregnancy | 0.98 | 0.016 | Mule deer | WY | High development | 2015 | Northrup et al. 2021 |
| Adult pregnancy | 0.94 | 0.05 | Mule deer | WY | Low development | 2009 | Northrup et al. 2021 |
| Adult pregnancy | 0.96 | 0.035 | Mule deer | WY | Low development | 2010 | Northrup et al. 2021 |
| Adult pregnancy | 0.98 | 0.02 | Mule deer | WY | Low development | 2011 | Northrup et al. 2021 |
| Adult pregnancy | 0.97 | 0.033 | Mule deer | WY | Low development | 2012 | Northrup et al. 2021 |
| Adult pregnancy | 0.93 | 0.045 | Mule deer | WY | Low development | 2013 | Northrup et al. 2021 |
| Adult pregnancy | 0.87 | 0.061 | Mule deer | WY | Low development | 2014 | Northrup et al. 2021 |
| Adult pregnancy | 0.96 | 0.037 | Mule deer | WY | Low development | 2015 | Northrup et al. 2021 |
| Adult pregnancy | 0.967 | 0.033 | Mule deer | OR | Murderers Creek | 2023 | ODFW unpublished |
| Adult pregnancy | 0.97 | 0.026 | Mule deer | SA |  | 2010 | Perera 2012 |
| Adult pregnancy | 0.97 | 0.03 | Mule deer | SA |  | 2011 | Perera 2012 |
| Adult pregnancy | 0.93 | 0.04 | Mule deer | CO |  | 1999 | Pojar and Bowden 2004/ Andelt et al. 2004 |
| Adult pregnancy | 0.95 | 0.02 | Mule deer | AZ |  | 2007-2008 | Quintana et al 2016 |
| Adult pregnancy | 0.913 | 0.02 | Mule deer | UT |  | 1948-49 | Robinette and Gashwiler 1950 |
| Adult pregnancy | 0.94 | 0.011 | Mule deer | UT |  | 1950-53 | Robinette et al. 1955 |
| Adult pregnancy | 0.97 | 0.027 | Mule deer | BC |  | 1997-2000 | Robinson et al. 2002 |
| Adult pregnancy | 0.98 | 0.02 | Mule deer | UT |  | 1979 | Smith 1983 |
| Adult pregnancy | 0.918 | 0.039 | Mule deer | UT |  | 1980 | Smith 1983 |
| Adult pregnancy | 0.96 | 0.03 | Mule deer | NM |  | 2011 | Sorensen 2015 |
| Adult pregnancy | 0.93 | 0.03 | Black-tailed deer | BC |  | 1963-67 | Thomas 1983 |
| Adult pregnancy | 0.95 | 0.06 | Mule deer | WA | East Slope Cascades | 2000-2007 | WDFW 2016 |
| Adult pregnancy | 0.92 | 0.1 | Mule deer | WA | Northern Rocky Mtns | 2000-2007 | WDFW 2016 |
| Adult pregnancy | 0.96 | 0.023 | Mule deer | WA | Columbia Plateau | 2000-2008 | WDFW 2021 |
| Adult pregnancy | 0.93 | 0.048 | Mule deer | WA | Okanagan Highlands | 2000-2007 | WDFW 2021 |
| Adult pregnancy | 0.949 | 0.016 | hybrid | CA | Overall | 2017-2020 | Wittmer et al. 2021 |
| Adult pregnancy | 0.954 | 0.031 | Mule deer | MT |  | 1983-85 | Wood 1987 |
| Juvenile survival (annual) | 0.047 | 0.045 | Mule deer | NM |  | 2002 | Bender et al. 2007/ Lomas and Bender 2007 |
| Juvenile survival (annual) | 0.08 | 0.08 | Mule deer | NM |  | 2003 | Bender et al. 2007/ Lomas and Bender 2007 |
| Juvenile survival (annual) | 0.26 | 0.04 | Black-tailed deer | CA |  | 2009-2012 | Forrester and Wittmer 2019; Marescot et al. 2015 |
| Juvenile survival (annual) | 0.39 | 0.11 | Black-tailed deer | AK |  | 2010 | Gilbert et al. 2020 |
| Juvenile survival (annual) | 0.084 | 0.09 | Black-tailed deer | AK |  | 2011 | Gilbert et al. 2020 |
| Juvenile survival (annual) | 0.52 | 0.127 | Black-tailed deer | AK |  | 2012 | Gilbert et al. 2020 |
| Juvenile survival (annual) | 0.453 | 0.077 | Mule deer | TX |  | 2004-2005 | Haskell et al. 2017 |
| Juvenile survival (annual) | 0.23 | 0.059 | Mule deer | TX |  | 2005-2006 | Haskell et al. 2017 |
| Juvenile survival (annual) | 0.19 | 0.079 | Mule deer | TX |  | 2006-2007 | Haskell et al. 2017 |
| Juvenile survival (annual) | 0.589 | 0.058 | Mule deer | CO |  | 2000 | Pojar and Bowden 2004 |
| Juvenile survival (annual) | 0.594 | 0.062 | Mule deer | CO |  | 2001 | Pojar and Bowden 2004 |
| Juvenile survival (annual) | 0.321 | 0.1 | Mule deer | CO |  | 1999 | Pojar and Bowden 2004/ Andelt et al. 2004 |
| Juvenile survival (annual) | 0.071 | 0.073 | Mule deer | AZ |  | 2007-2008 | Quintana et al 2016 |
| Juvenile survival (annual) | 0.49 | 0.08 | Mule deer | SA |  | 2007 | Skelton 2010 |
| Juvenile survival (annual) | 0.34 | 0.06 | Mule deer | OR |  | 2010-2012 | Speten 2014 |
| Juvenile survival (annual) | 0.27 | 0.05 | hybrid | CA |  | 2017-2018 | Wittmer et al. 2021 |
| Juvenile survival (annual) | 0.217 | 0.063 | hybrid | CA |  | 2018-2019 | Wittmer et al. 2021 |
| Juvenile survival (annual) | 0.461 | 0.072 | hybrid | CA |  | 2019-2020 | Wittmer et al. 2021 |
| Juvenile survival (summer) | 0.047 | 0.045 | Mule deer | NM |  | 2002 | Bender et al. 2007/ Lomas and Bender 2007 |
| Juvenile survival (summer) | 0.12 | 0.06 | Mule deer | NM |  | 2003 | Bender et al. 2007/ Lomas and Bender 2007 |
| Juvenile survival (summer) | 0.52 | 0.07 | Mule deer | NM |  | 2004 | Bender et al. 2007/ Lomas and Bender 2007 |
| Juvenile survival (summer) | 0.436 | 0.129 | Mule deer | CO | Control | 2001-2002 | Bishop et al. 2009 |
| Juvenile survival (summer) | 0.488 | 0.079 | Mule deer | CO | Control | 2002-2003 | Bishop et al. 2009 |
| Juvenile survival (summer) | 0.336 | 0.075 | Mule deer | CO | Control | 2003-2004 | Bishop et al. 2009 |
| Juvenile survival (summer) | 0.527 | 0.122 | Mule deer | CO | Treatment | 2001-2002 | Bishop et al. 2009 |
| Juvenile survival (summer) | 0.553 | 0.076 | Mule deer | CO | Treatment | 2002-2003 | Bishop et al. 2009 |
| Juvenile survival (summer) | 0.47 | 0.083 | Mule deer | CO | Treatment | 2003-2004 | Bishop et al. 2009 |
| Juvenile survival (summer) | 0.226 | 0.076 | Mule deer | CA |  | 1993 | Bleich et al. 2006 |
| Juvenile survival (summer) | 0.228 | 0.083 | Mule deer | CA |  | 1994 | Bleich et al. 2006 |
| Juvenile survival (summer) | 0.37 | 0.081 | Mule deer | CA |  | 1995 | Bleich et al. 2006 |
| Juvenile survival (summer) | 0.292 | 0.098 | Mule deer | CA |  | 1996 | Bleich et al. 2006 |
| Juvenile survival (summer) | 0.3 | 0.08 | Black-tailed deer | BC | Adam | 1980-81 | Hatter 1988 |
| Juvenile survival (summer) | 0.376 | 0.07 | Mule deer | ID | Overall | 1998 | Hurley et al. 2011 |
| Juvenile survival (summer) | 0.533 | 0.039 | Mule deer | ID | Overall | 1999 | Hurley et al. 2011 |
| Juvenile survival (summer) | 0.475 | 0.115 | Mule deer | ID | Reference | 1998 | Hurley et al. 2011 |
| Juvenile survival (summer) | 0.377 | 0.056 | Mule deer | ID | Reference | 1999 | Hurley et al. 2011 |
| Juvenile survival (summer) | 0.266 | 0.048 | Mule deer | ID | Reference | 2000 | Hurley et al. 2011 |
| Juvenile survival (summer) | 0.607 | 0.046 | Mule deer | ID | Reference | 2001 | Hurley et al. 2011 |
| Juvenile survival (summer) | 0.488 | 0.049 | Mule deer | ID | Reference | 2002 | Hurley et al. 2011 |
| Juvenile survival (summer) | 0.277 | 0.065 | Mule deer | ID | Treatment | 1998 | Hurley et al. 2011 |
| Juvenile survival (summer) | 0.689 | 0.049 | Mule deer | ID | Treatment | 1999 | Hurley et al. 2011 |
| Juvenile survival (summer) | 0.196 | 0.056 | Mule deer | ID | Treatment | 2000 | Hurley et al. 2011 |
| Juvenile survival (summer) | 0.609 | 0.052 | Mule deer | ID | Treatment | 2001 | Hurley et al. 2011 |
| Juvenile survival (summer) | 0.74 | 0.045 | Mule deer | ID | Treatment | 2002 | Hurley et al. 2011 |
| Juvenile survival (summer) | 0.59 | 0.053 | Black-tailed deer | WA |  | 2006 | McCoy and Murphie 2011/McCoy et al. 2014 |
| Juvenile survival (summer) | 0.57 | 0.053 | Black-tailed deer | WA |  | 2007 | McCoy and Murphie 2011/McCoy et al. 2014 |
| Juvenile survival (summer) | 0.58 | 0.053 | Black-tailed deer | WA |  | 2008 | McCoy and Murphie 2011/McCoy et al. 2014 |
| Juvenile survival (summer) | 0.39 | 0.053 | Black-tailed deer | WA |  | 2009 | McCoy and Murphie 2011/McCoy et al. 2014 |
| Juvenile survival (summer) | 0.347 | 0.083 | Mule deer | SA |  | 2009 | Perera 2012 |
| Juvenile survival (summer) | 0.335 | 0.078 | Mule deer | SA |  | 2010 | Perera 2012 |
| Juvenile survival (summer) | 0.34 | 0.083 | Mule deer | SA |  | 2011 | Perera 2012 |
| Juvenile survival (summer) | 0.496 | 0.076 | Mule deer | UT |  | 1979 | Smith 1983 |
| Juvenile survival (summer) | 0.358 | 0.09 | Mule deer | UT |  | 1980 | Smith 1983 |
| Juvenile survival (summer) | 0.459 | 0.074 | hybrid | CA | Overall | 2017-2020 | Wittmer et al. 2021 |
| Juvenile survival (winter) | 0.675 | 0.112 | Mule deer | CO | High treatment | 2005-2008 | Bergman et al. 2014 |
| Juvenile survival (winter) | 0.768 | 0.085 | Mule deer | CO | Low/no treatment | 2005-2008 | Bergman et al. 2014 |
| Juvenile survival (winter) | 0.063 | 0.06 | Mule deer | ID | Bennett | 1992-93 | Bishop et al. 2005 |
| Juvenile survival (winter) | 0.857 | 0.076 | Mule deer | ID | Bennett | 1993-94 | Bishop et al. 2005 |
| Juvenile survival (winter) | 0.263 | 0.101 | Mule deer | ID | Bennett | 1994-95 | Bishop et al. 2005 |
| Juvenile survival (winter) | 0.524 | 0.109 | Mule deer | ID | Bennett | 1995-96 | Bishop et al. 2005 |
| Juvenile survival (winter) | 0.406 | 0.104 | Mule deer | ID | Bennett | 1996-97 | Bishop et al. 2005 |
| Juvenile survival (winter) | 0.238 | 0.093 | Mule deer | ID | Blacks Creek | 1992-93 | Bishop et al. 2005 |
| Juvenile survival (winter) | 0.9 | 0.069 | Mule deer | ID | Blacks Creek | 1993-94 | Bishop et al. 2005 |
| Juvenile survival (winter) | 0.7 | 0.102 | Mule deer | ID | Blacks Creek | 1994-95 | Bishop et al. 2005 |
| Juvenile survival (winter) | 0.842 | 0.084 | Mule deer | ID | Blacks Creek | 1995-96 | Bishop et al. 2005 |
| Juvenile survival (winter) | 0.431 | 0.115 | Mule deer | ID | Blacks Creek | 1996-97 | Bishop et al. 2005 |
| Juvenile survival (winter) | 0.474 | 0.14 | Mule deer | ID | Owyhee | 1992-93 | Bishop et al. 2005 |
| Juvenile survival (winter) | 0.474 | 0.13 | Mule deer | ID | Owyhee | 1993-94 | Bishop et al. 2005 |
| Juvenile survival (winter) | 0.4 | 0.117 | Mule deer | ID | Owyhee | 1994-95 | Bishop et al. 2005 |
| Juvenile survival (winter) | 0.905 | 0.064 | Mule deer | ID | Owyhee | 1995-96 | Bishop et al. 2005 |
| Juvenile survival (winter) | 0.476 | 0.115 | Mule deer | ID | Owyhee | 1996-97 | Bishop et al. 2005 |
| Juvenile survival (winter) | 0.648 | 0.081 | Mule deer | CO | Control | 2001-2002 | Bishop et al. 2009 |
| Juvenile survival (winter) | 0.763 | 0.069 | Mule deer | CO | Control | 2002-2003 | Bishop et al. 2009 |
| Juvenile survival (winter) | 0.78 | 0.064 | Mule deer | CO | Control | 2003-2004 | Bishop et al. 2009 |
| Juvenile survival (winter) | 0.894 | 0.038 | Mule deer | CO | Treatment | 2001-2002 | Bishop et al. 2009 |
| Juvenile survival (winter) | 0.932 | 0.027 | Mule deer | CO | Treatment | 2002-2003 | Bishop et al. 2009 |
| Juvenile survival (winter) | 0.938 | 0.025 | Mule deer | CO | Treatment | 2003-2004 | Bishop et al. 2009 |
| Juvenile survival (winter) | 0.786 | 0.012 | Mule deer | CA |  | 1993 | Bleich et al. 2006 |
| Juvenile survival (winter) | 0.933 | 0.004 | Mule deer | CA |  | 1994 | Bleich et al. 2006 |
| Juvenile survival (winter) | 0.882 | 0.006 | Mule deer | CA |  | 1995 | Bleich et al. 2006 |
| Juvenile survival (winter) | 0.833 | 0.012 | Mule deer | CA |  | 1996 | Bleich et al. 2006 |
| Juvenile survival (winter) | 0.65 | 0.039 | Mule deer | ID | Overall | 1998 | Hurley et al. 2011 |
| Juvenile survival (winter) | 0.689 | 0.032 | Mule deer | ID | Overall | 1999 | Hurley et al. 2011 |
| Juvenile survival (winter) | 0.59 | 0.056 | Mule deer | ID | Reference | 1998 | Hurley et al. 2011 |
| Juvenile survival (winter) | 0.644 | 0.047 | Mule deer | ID | Reference | 1999 | Hurley et al. 2011 |
| Juvenile survival (winter) | 0.507 | 0.048 | Mule deer | ID | Reference | 2000 | Hurley et al. 2011 |
| Juvenile survival (winter) | 0.726 | 0.045 | Mule deer | ID | Reference | 2001 | Hurley et al. 2011 |
| Juvenile survival (winter) | 0.308 | 0.043 | Mule deer | ID | Reference | 2002 | Hurley et al. 2011 |
| Juvenile survival (winter) | 0.703 | 0.054 | Mule deer | ID | Treatment | 1998 | Hurley et al. 2011 |
| Juvenile survival (winter) | 0.733 | 0.043 | Mule deer | ID | Treatment | 1999 | Hurley et al. 2011 |
| Juvenile survival (winter) | 0.828 | 0.036 | Mule deer | ID | Treatment | 2000 | Hurley et al. 2011 |
| Juvenile survival (winter) | 0.724 | 0.044 | Mule deer | ID | Treatment | 2001 | Hurley et al. 2011 |
| Juvenile survival (winter) | 0.081 | 0.029 | Mule deer | ID | Treatment | 2002 | Hurley et al. 2011 |
| Juvenile survival (winter) | 0.48 | 0.099 | Mule deer | ID | Bannock | 2004 | Hurley et al. 2017 |
| Juvenile survival (winter) | 0.74 | 0.092 | Mule deer | ID | Bannock | 2007 | Hurley et al. 2017 |
| Juvenile survival (winter) | 0.33 | 0.093 | Mule deer | ID | Bannock | 2008 | Hurley et al. 2017 |
| Juvenile survival (winter) | 0.36 | 0.071 | Mule deer | ID | Bannock | 2009 | Hurley et al. 2017 |
| Juvenile survival (winter) | 0.71 | 0.082 | Mule deer | ID | Bannock | 2010 | Hurley et al. 2017 |
| Juvenile survival (winter) | 0.56 | 0.099 | Mule deer | ID | Boise River | 2003 | Hurley et al. 2017 |
| Juvenile survival (winter) | 0.35 | 0.099 | Mule deer | ID | Boise River | 2004 | Hurley et al. 2017 |
| Juvenile survival (winter) | 0.74 | 0.092 | Mule deer | ID | Boise River | 2005 | Hurley et al. 2017 |
| Juvenile survival (winter) | 0.52 | 0.1 | Mule deer | ID | Boise River | 2006 | Hurley et al. 2017 |
| Juvenile survival (winter) | 0.46 | 0.103 | Mule deer | ID | Boise River | 2007 | Hurley et al. 2017 |
| Juvenile survival (winter) | 0.71 | 0.104 | Mule deer | ID | Boise River | 2008 | Hurley et al. 2017 |
| Juvenile survival (winter) | 0.77 | 0.092 | Mule deer | ID | Boise River | 2009 | Hurley et al. 2017 |
| Juvenile survival (winter) | 0.76 | 0.084 | Mule deer | ID | Boise River | 2010 | Hurley et al. 2017 |
| Juvenile survival (winter) | 0.48 | 0.099 | Mule deer | ID | Boise River | 2011 | Hurley et al. 2017 |
| Juvenile survival (winter) | 0.67 | 0.096 | Mule deer | ID | Boise River | 2012 | Hurley et al. 2017 |
| Juvenile survival (winter) | 0.7 | 0.096 | Mule deer | ID | Boise River | 2013 | Hurley et al. 2017 |
| Juvenile survival (winter) | 0.35 | 0.099 | Mule deer | ID | C. Mountains | 2003 | Hurley et al. 2017 |
| Juvenile survival (winter) | 0.32 | 0.099 | Mule deer | ID | C. Mountains | 2004 | Hurley et al. 2017 |
| Juvenile survival (winter) | 0.67 | 0.086 | Mule deer | ID | C. Mountains | 2005 | Hurley et al. 2017 |
| Juvenile survival (winter) | 0.1 | 0.044 | Mule deer | ID | C. Mountains | 2006 | Hurley et al. 2017 |
| Juvenile survival (winter) | 0.64 | 0.059 | Mule deer | ID | C. Mountains | 2007 | Hurley et al. 2017 |
| Juvenile survival (winter) | 0.42 | 0.114 | Mule deer | ID | C. Mountains | 2008 | Hurley et al. 2017 |
| Juvenile survival (winter) | 0.39 | 0.092 | Mule deer | ID | C. Mountains | 2009 | Hurley et al. 2017 |
| Juvenile survival (winter) | 0.87 | 0.06 | Mule deer | ID | C. Mountains | 2010 | Hurley et al. 2017 |
| Juvenile survival (winter) | 0.47 | 0.076 | Mule deer | ID | C. Mountains | 2011 | Hurley et al. 2017 |
| Juvenile survival (winter) | 0.61 | 0.085 | Mule deer | ID | C. Mountains | 2012 | Hurley et al. 2017 |
| Juvenile survival (winter) | 0.47 | 0.091 | Mule deer | ID | C. Mountains | 2013 | Hurley et al. 2017 |
| Juvenile survival (winter) | 0.74 | 0.063 | Mule deer | ID | Caribou | 2003 | Hurley et al. 2017 |
| Juvenile survival (winter) | 0.53 | 0.106 | Mule deer | ID | Caribou | 2004 | Hurley et al. 2017 |
| Juvenile survival (winter) | 0.52 | 0.099 | Mule deer | ID | Caribou | 2005 | Hurley et al. 2017 |
| Juvenile survival (winter) | 0.31 | 0.055 | Mule deer | ID | Caribou | 2006 | Hurley et al. 2017 |
| Juvenile survival (winter) | 0.81 | 0.054 | Mule deer | ID | Caribou | 2007 | Hurley et al. 2017 |
| Juvenile survival (winter) | 0.22 | 0.072 | Mule deer | ID | Caribou | 2008 | Hurley et al. 2017 |
| Juvenile survival (winter) | 0.28 | 0.076 | Mule deer | ID | Caribou | 2009 | Hurley et al. 2017 |
| Juvenile survival (winter) | 0.61 | 0.092 | Mule deer | ID | Caribou | 2010 | Hurley et al. 2017 |
| Juvenile survival (winter) | 0.85 | 0.071 | Mule deer | ID | Island Park | 2004 | Hurley et al. 2017 |
| Juvenile survival (winter) | 0.32 | 0.093 | Mule deer | ID | Island Park | 2008 | Hurley et al. 2017 |
| Juvenile survival (winter) | 0.56 | 0.111 | Mule deer | ID | Island Park | 2009 | Hurley et al. 2017 |
| Juvenile survival (winter) | 0.68 | 0.109 | Mule deer | ID | Island Park | 2010 | Hurley et al. 2017 |
| Juvenile survival (winter) | 0.07 | 0.057 | Mule deer | ID | Island Park | 2011 | Hurley et al. 2017 |
| Juvenile survival (winter) | 0.24 | 0.103 | Mule deer | ID | Middle Fork | 2008 | Hurley et al. 2017 |
| Juvenile survival (winter) | 0.57 | 0.103 | Mule deer | ID | Mountain Valley | 2004 | Hurley et al. 2017 |
| Juvenile survival (winter) | 0.89 | 0.047 | Mule deer | ID | Mountain Valley | 2005 | Hurley et al. 2017 |
| Juvenile survival (winter) | 0.17 | 0.076 | Mule deer | ID | Mountain Valley | 2006 | Hurley et al. 2017 |
| Juvenile survival (winter) | 0.63 | 0.112 | Mule deer | ID | Mountain Valley | 2007 | Hurley et al. 2017 |
| Juvenile survival (winter) | 0.35 | 0.112 | Mule deer | ID | Mountain Valley | 2008 | Hurley et al. 2017 |
| Juvenile survival (winter) | 0.39 | 0.106 | Mule deer | ID | Mountain Valley | 2009 | Hurley et al. 2017 |
| Juvenile survival (winter) | 0.69 | 0.075 | Mule deer | ID | Mountain Valley | 2010 | Hurley et al. 2017 |
| Juvenile survival (winter) | 0.22 | 0.076 | Mule deer | ID | Mountain Valley | 2011 | Hurley et al. 2017 |
| Juvenile survival (winter) | 0.39 | 0.083 | Mule deer | ID | Mountain Valley | 2012 | Hurley et al. 2017 |
| Juvenile survival (winter) | 0.37 | 0.08 | Mule deer | ID | Mountain Valley | 2013 | Hurley et al. 2017 |
| Juvenile survival (winter) | 0.92 | 0.054 | Mule deer | ID | Palisades | 2003 | Hurley et al. 2017 |
| Juvenile survival (winter) | 0.54 | 0.098 | Mule deer | ID | Palisades | 2004 | Hurley et al. 2017 |
| Juvenile survival (winter) | 0.68 | 0.094 | Mule deer | ID | Palisades | 2005 | Hurley et al. 2017 |
| Juvenile survival (winter) | 0.16 | 0.073 | Mule deer | ID | Palisades | 2006 | Hurley et al. 2017 |
| Juvenile survival (winter) | 0.64 | 0.096 | Mule deer | ID | Palisades | 2007 | Hurley et al. 2017 |
| Juvenile survival (winter) | 0.09 | 0.087 | Mule deer | ID | Palisades | 2008 | Hurley et al. 2017 |
| Juvenile survival (winter) | 0.52 | 0.109 | Mule deer | ID | Palisades | 2009 | Hurley et al. 2017 |
| Juvenile survival (winter) | 0.75 | 0.097 | Mule deer | ID | Palisades | 2010 | Hurley et al. 2017 |
| Juvenile survival (winter) | 0.32 | 0.099 | Mule deer | ID | Smokey-Bennett | 2008 | Hurley et al. 2017 |
| Juvenile survival (winter) | 0.67 | 0.09 | Mule deer | ID | Smokey-Bennett | 2009 | Hurley et al. 2017 |
| Juvenile survival (winter) | 0.83 | 0.076 | Mule deer | ID | Smokey-Bennett | 2010 | Hurley et al. 2017 |
| Juvenile survival (winter) | 0.37 | 0.093 | Mule deer | ID | Smokey-Bennett | 2011 | Hurley et al. 2017 |
| Juvenile survival (winter) | 0.82 | 0.067 | Mule deer | ID | Smokey-Bennett | 2012 | Hurley et al. 2017 |
| Juvenile survival (winter) | 0.85 | 0.063 | Mule deer | ID | Smokey-Bennett | 2013 | Hurley et al. 2017 |
| Juvenile survival (winter) | 0.75 | 0.089 | Mule deer | ID | South Hills | 2003 | Hurley et al. 2017 |
| Juvenile survival (winter) | 0.83 | 0.079 | Mule deer | ID | South Hills | 2004 | Hurley et al. 2017 |
| Juvenile survival (winter) | 0.73 | 0.087 | Mule deer | ID | South Hills | 2005 | Hurley et al. 2017 |
| Juvenile survival (winter) | 0.32 | 0.105 | Mule deer | ID | South Hills | 2006 | Hurley et al. 2017 |
| Juvenile survival (winter) | 0.57 | 0.126 | Mule deer | ID | South Hills | 2007 | Hurley et al. 2017 |
| Juvenile survival (winter) | 0.35 | 0.107 | Mule deer | ID | South Hills | 2008 | Hurley et al. 2017 |
| Juvenile survival (winter) | 0.3 | 0.113 | Mule deer | ID | South Hills | 2009 | Hurley et al. 2017 |
| Juvenile survival (winter) | 0.85 | 0.071 | Mule deer | ID | South Hills | 2010 | Hurley et al. 2017 |
| Juvenile survival (winter) | 0.65 | 0.101 | Mule deer | ID | South Hills | 2012 | Hurley et al. 2017 |
| Juvenile survival (winter) | 0.59 | 0.113 | Mule deer | ID | South Hills | 2013 | Hurley et al. 2017 |
| Juvenile survival (winter) | 0.64 | 0.096 | Mule deer | ID | Weiser-McCall | 2003 | Hurley et al. 2017 |
| Juvenile survival (winter) | 0.41 | 0.07 | Mule deer | ID | Weiser-McCall | 2004 | Hurley et al. 2017 |
| Juvenile survival (winter) | 0.95 | 0.051 | Mule deer | ID | Weiser-McCall | 2005 | Hurley et al. 2017 |
| Juvenile survival (winter) | 0.43 | 0.094 | Mule deer | ID | Weiser-McCall | 2006 | Hurley et al. 2017 |
| Juvenile survival (winter) | 0.67 | 0.111 | Mule deer | ID | Weiser-McCall | 2007 | Hurley et al. 2017 |
| Juvenile survival (winter) | 0.32 | 0.101 | Mule deer | ID | Weiser-McCall | 2008 | Hurley et al. 2017 |
| Juvenile survival (winter) | 0.86 | 0.074 | Mule deer | ID | Weiser-McCall | 2009 | Hurley et al. 2017 |
| Juvenile survival (winter) | 0.55 | 0.084 | Mule deer | ID | Weiser-McCall | 2010 | Hurley et al. 2017 |
| Juvenile survival (winter) | 0.09 | 0.052 | Mule deer | ID | Weiser-McCall | 2011 | Hurley et al. 2017 |
| Juvenile survival (winter) | 0.67 | 0.086 | Mule deer | ID | Weiser-McCall | 2012 | Hurley et al. 2017 |
| Juvenile survival (winter) | 0.69 | 0.082 | Mule deer | ID | Weiser-McCall | 2013 | Hurley et al. 2017 |
| Juvenile survival (winter) | 0.77 | 0.118 | Mule deer | MT |  | 2014-2016 | Kolar et al. 2018 |
| Juvenile survival (winter) | 0.692 | 0.062 | Mule deer | CO | D-16 | 1999–2000 | Lukacs et al. 2009 |
| Juvenile survival (winter) | 0.529 | 0.065 | Mule deer | CO | D-16 | 2000–2001 | Lukacs et al. 2009 |
| Juvenile survival (winter) | 0.703 | 0.064 | Mule deer | CO | D-16 | 2001–2002 | Lukacs et al. 2009 |
| Juvenile survival (winter) | 0.674 | 0.069 | Mule deer | CO | D-16 | 2002–2003 | Lukacs et al. 2009 |
| Juvenile survival (winter) | 0.868 | 0.044 | Mule deer | CO | D-16 | 2003–2004 | Lukacs et al. 2009 |
| Juvenile survival (winter) | 0.833 | 0.048 | Mule deer | CO | D-16 | 2004–2005 | Lukacs et al. 2009 |
| Juvenile survival (winter) | 0.815 | 0.05 | Mule deer | CO | D-16 | 2005–2006 | Lukacs et al. 2009 |
| Juvenile survival (winter) | 0.843 | 0.048 | Mule deer | CO | D-16 | 2006–2007 | Lukacs et al. 2009 |
| Juvenile survival (winter) | 0.898 | 0.039 | Mule deer | CO | D-16 | 2007–2008 | Lukacs et al. 2009 |
| Juvenile survival (winter) | 0.519 | 0.079 | Mule deer | CO | D-19 | 1997–1998 | Lukacs et al. 2009 |
| Juvenile survival (winter) | 0.709 | 0.061 | Mule deer | CO | D-19 | 1998–1999 | Lukacs et al. 2009 |
| Juvenile survival (winter) | 0.62 | 0.063 | Mule deer | CO | D-19 | 1999–2000 | Lukacs et al. 2009 |
| Juvenile survival (winter) | 0.702 | 0.064 | Mule deer | CO | D-19 | 2000–2001 | Lukacs et al. 2009 |
| Juvenile survival (winter) | 0.482 | 0.095 | Mule deer | CO | D-19 | 2001–2002 | Lukacs et al. 2009 |
| Juvenile survival (winter) | 0.892 | 0.045 | Mule deer | CO | D-19 | 2002–2003 | Lukacs et al. 2009 |
| Juvenile survival (winter) | 0.839 | 0.052 | Mule deer | CO | D-19 | 2003–2004 | Lukacs et al. 2009 |
| Juvenile survival (winter) | 0.865 | 0.047 | Mule deer | CO | D-19 | 2004–2005 | Lukacs et al. 2009 |
| Juvenile survival (winter) | 0.918 | 0.035 | Mule deer | CO | D-19 | 2005–2006 | Lukacs et al. 2009 |
| Juvenile survival (winter) | 0.762 | 0.054 | Mule deer | CO | D-19 | 2006–2007 | Lukacs et al. 2009 |
| Juvenile survival (winter) | 0.739 | 0.058 | Mule deer | CO | D-19 | 2007–2008 | Lukacs et al. 2009 |
| Juvenile survival (winter) | 0.725 | 0.065 | Mule deer | CO | D-7 | 2001–2002 | Lukacs et al. 2009 |
| Juvenile survival (winter) | 0.801 | 0.059 | Mule deer | CO | D-7 | 2002–2003 | Lukacs et al. 2009 |
| Juvenile survival (winter) | 0.651 | 0.07 | Mule deer | CO | D-7 | 2003–2004 | Lukacs et al. 2009 |
| Juvenile survival (winter) | 0.823 | 0.054 | Mule deer | CO | D-7 | 2004–2005 | Lukacs et al. 2009 |
| Juvenile survival (winter) | 0.636 | 0.068 | Mule deer | CO | D-7 | 2005–2006 | Lukacs et al. 2009 |
| Juvenile survival (winter) | 0.753 | 0.057 | Mule deer | CO | D-7 | 2006–2007 | Lukacs et al. 2009 |
| Juvenile survival (winter) | 0.34 | 0.059 | Mule deer | CO | D-7 | 2007–2008 | Lukacs et al. 2009 |
| Juvenile survival (winter) | 0.866 | 0.051 | Mule deer | CO | D-9 | 1998–1999 | Lukacs et al. 2009 |
| Juvenile survival (winter) | 0.767 | 0.059 | Mule deer | CO | D-9 | 1999–2000 | Lukacs et al. 2009 |
| Juvenile survival (winter) | 0.806 | 0.055 | Mule deer | CO | D-9 | 2000–2001 | Lukacs et al. 2009 |
| Juvenile survival (winter) | 0.721 | 0.059 | Mule deer | CO | D-9 | 2001–2002 | Lukacs et al. 2009 |
| Juvenile survival (winter) | 0.82 | 0.049 | Mule deer | CO | D-9 | 2002–2003 | Lukacs et al. 2009 |
| Juvenile survival (winter) | 0.9 | 0.039 | Mule deer | CO | D-9 | 2003–2004 | Lukacs et al. 2009 |
| Juvenile survival (winter) | 0.61 | 0.063 | Mule deer | CO | D-9 | 2004–2005 | Lukacs et al. 2009 |
| Juvenile survival (winter) | 0.51 | 0.065 | Mule deer | CO | D-9 | 2005–2006 | Lukacs et al. 2009 |
| Juvenile survival (winter) | 0.615 | 0.062 | Mule deer | CO | D-9 | 2006–2007 | Lukacs et al. 2009 |
| Juvenile survival (winter) | 0.412 | 0.062 | Mule deer | CO | D-9 | 2007–2008 | Lukacs et al. 2009 |
| Juvenile survival (winter) | 0.56 | 0.05 | Black-tailed deer | WA |  | 2006 | McCoy and Murphie 2011/McCoy et al. 2014 |
| Juvenile survival (winter) | 0.67 | 0.05 | Black-tailed deer | WA |  | 2007 | McCoy and Murphie 2011/McCoy et al. 2014 |
| Juvenile survival (winter) | 0.67 | 0.05 | Black-tailed deer | WA |  | 2008 | McCoy and Murphie 2011/McCoy et al. 2014 |
| Juvenile survival (winter) | 0.78 | 0.05 | Black-tailed deer | WA |  | 2009 | McCoy and Murphie 2011/McCoy et al. 2014 |
| Juvenile survival (winter) | 0.583 | 0.074 | Mule deer | UT |  | 1979 | Smith 1983 |
| Juvenile survival (winter) | 0.889 | 0.018 | Mule deer | UT |  | 1980 | Smith 1983 |
| Juvenile survival (winter) | 0.525 | 0.096 | Mule deer | MT |  | 1989 | Unsworth et al. 1999 |
| Juvenile survival (winter) | 0.68 | 0.072 | Mule deer | MT |  | 1990 | Unsworth et al. 1999 |
| Juvenile survival (winter) | 0.6 | 0.083 | Mule deer | MT |  | 1991 | Unsworth et al. 1999 |
| Juvenile survival (winter) | 0.233 | 0.074 | Mule deer | MT |  | 1992 | Unsworth et al. 1999 |
| Juvenile survival (winter) | 0.229 | 0.058 | Mule deer | ID |  | 1992 | Unsworth et al. 1999 |
| Juvenile survival (winter) | 0.625 | 0.084 | Mule deer | MT |  | 1993 | Unsworth et al. 1999 |
| Juvenile survival (winter) | 0.76 | 0.057 | Mule deer | ID |  | 1993 | Unsworth et al. 1999 |
| Juvenile survival (winter) | 0.137 | 0.051 | Mule deer | MT |  | 1994 | Unsworth et al. 1999 |
| Juvenile survival (winter) | 0.469 | 0.067 | Mule deer | ID |  | 1994 | Unsworth et al. 1999 |
| Juvenile survival (winter) | 0.759 | 0.055 | Mule deer | ID |  | 1995 | Unsworth et al. 1999 |
| Juvenile survival (winter) | 0.448 | 0.065 | Mule deer | ID |  | 1996 | Unsworth et al. 1999 |
| Juvenile survival (winter) | 0.32 | 0.09 | Mule deer | CO |  | 1982 | White and Bartmann 1998 |
| Juvenile survival (winter) | 0.07 | 0.05 | Mule deer | CO |  | 1983 | White and Bartmann 1998 |
| Juvenile survival (winter) | 0.2 | 0.08 | Mule deer | CO |  | 1984 | White and Bartmann 1998 |
| Juvenile survival (winter) | 0.54 | 0.07 | Mule deer | CO |  | 1985 | White and Bartmann 1998 |
| Juvenile survival (winter) | 0.43 | 0.06 | Mule deer | CO |  | 1986 | White and Bartmann 1998 |
| Juvenile survival (winter) | 0.24 | 0.08 | Mule deer | CO |  | 1987 | White and Bartmann 1998 |
| Juvenile survival (winter) | 0.27 | 0.08 | Mule deer | CO |  | 1988 | White and Bartmann 1998 |
| Juvenile survival (winter) | 0.78 | 0.08 | Mule deer | CO |  | 1989 | White and Bartmann 1998 |
| Juvenile survival (winter) | 0.32 | 0.09 | Mule deer | CO |  | 1990 | White and Bartmann 1998 |
| Juvenile survival (winter) | 0.46 | 0.11 | Mule deer | CO |  | 1991 | White and Bartmann 1998 |
| Juvenile survival (winter) | 0.11 | 0.04 | Mule deer | CO |  | 1992 | White and Bartmann 1998 |
| Juvenile survival (winter) | 0.55 | 0.06 | Mule deer | CO |  | 1993 | White and Bartmann 1998 |
| Juvenile survival (winter) | 0.7 | 0.05 | Mule deer | CO |  | 1994 | White and Bartmann 1998 |
| Juvenile survival (winter) | 0.62 | 0.06 | Mule deer | CO |  | 1995 | White and Bartmann 1998 |
| Juvenile survival (winter) | 0.687 | 0.109 | hybrid | CA | Overall | 2017-2020 | Wittmer et al. 2021 |
| Yearling female survival (annual) | 0.825 | 0.054 | Mule deer | CO | Overall | 2002-2004 | Bishop et al. 2009 |
| Yearling female survival (annual) | 0.808 | 0.016 | Mule deer | CA |  | 1993 | Bleich et al. 2006 |
| Yearling female survival (annual) | 0.933 | 0.007 | Mule deer | CA |  | 1994 | Bleich et al. 2006 |
| Yearling female survival (annual) | 0.867 | 0.011 | Mule deer | CA |  | 1995 | Bleich et al. 2006 |
| Yearling female survival (annual) | 0.85 | 0.12 | Mule deer | CA |  | 1996 | Bleich et al. 2006 |
| Yearling female survival (annual) | 0.56 | 0.18 | Black-tailed deer | CA |  | 2009-2012 | Forrester and Wittmer 2019; Marescot et al. 2015 |
| Yearling female survival (annual) | 0.771 | 0.072 | Mule deer | WY |  | 1993-95 | Gogan et al. 2019 |
| Yearling female survival (annual) | 0.97 | 0.018 | Mule deer | CA | Overall | 1997-2009 | Monteith et al. 2014 |
| Yearling female survival (annual) | 0.799 | 0.07 | Mule deer | MT |  | 1953-63 | Nellis 1968 and Nellis 1964 |
| Yearling female survival (annual) | 0.934 | 0.07 | Black-tailed deer | CA | Chaparral | 1948-55 | Taber and Dasmann 1957 |
| Yearling female survival (annual) | 0.825 | 0.07 | Black-tailed deer | CA | Shrubland | 1948-55 | Taber and Dasmann 1957 |
| Yearling female survival (annual) | 0.741 | 0.161 | hybrid | CA | Overall | 2017-2020 | Wittmer et al. 2021 |
| Yearling litter size | 1.16 | 0.14 | Mule deer | NM |  | 2003-2008 | Bender and Hoenes 2018 |
| Yearling litter size | 1 | 0.19 | Mule deer | CO | Control | 2002-2003 | Bishop et al. 2009 |
| Yearling litter size | 1.053 | 0.2 | Mule deer | CO | Control | 2003-2004 | Bishop et al. 2009 |
| Yearling litter size | 1.034 | 0.19 | Mule deer | CO | Treatment | 2002-2003 | Bishop et al. 2009 |
| Yearling litter size | 1.386 | 0.2 | Mule deer | CO | Treatment | 2003-2004 | Bishop et al. 2009 |
| Yearling litter size | 1.33 | 0.24 | Black-tailed deer | BC | Adam | 1980-81 | Hatter 1988 |
| Yearling litter size | 1.14 | 0.12 | Black-tailed deer | BC | Nimpkish | 1980-81 | Hatter 1988 |
| Yearling litter size | 1.07 | 0.06 | Black-tailed deer | BC | Northwest | 1980-81 | Hatter 1988 |
| Yearling litter size | 1.14 | 0.12 | Mule deer | UT | Antimony | 1954-56 | Julander et al. 1961 |
| Yearling litter size | 1.56 | 0.17 | Mule deer | ID | Sublett | 1954-56 | Julander et al. 1961 |
| Yearling litter size | 1.37 | 0.11 | Mule deer | MT |  | 1953-63 | Nellis 1968 and Nellis 1964 |
| Yearling litter size | 1.83 | 0.15 | Mule deer | SA |  | 2010 | Perera 2012 |
| Yearling litter size | 1.92 | 0.078 | Mule deer | SA |  | 2011 | Perera 2012 |
| Yearling litter size | 1.375 | 0.086 | Mule deer | UT |  | 1948-49 | Robinette and Gashwiler 1950 |
| Yearling litter size | 1.32 | 0.037 | Mule deer | UT |  | 1950-53 | Robinette et al. 1955 |
| Yearling litter size | 1.5 | 0.158 | Mule deer | UT |  | 1979 | Smith 1983 |
| Yearling litter size | 1.14 | 0.127 | Mule deer | UT |  | 1980 | Smith 1983 |
| Yearling litter size | 1.09 | 0.086 | Black-tailed deer | BC |  | 1963-67 | Thomas 1983 |
| Yearling pregnancy | 0.4 | 0.17 | Mule deer | NM |  | 2003-2008 | Bender and Hoenes 2018 |
| Yearling pregnancy | 0.43 | 0.187 | Mule deer | MT | Cabinet-Salish | 2017-2019 | DeCesare et al. 2021 |
| Yearling pregnancy | 0.75 | 0.19 | Mule deer | MT | Rocky Mountain Front | 2017-2019 | DeCesare et al. 2021 |
| Yearling pregnancy | 0.82 | 0.115 | Mule deer | MT | Whitefish | 2017-2019 | DeCesare et al. 2021 |
| Yearling pregnancy | 0.67 | 0.24 | Black-tailed deer | BC | Adam | 1980-81 | Hatter 1988 |
| Yearling pregnancy | 0.6 | 0.09 | Black-tailed deer | BC | Nimpkish | 1980-81 | Hatter 1988 |
| Yearling pregnancy | 0.92 | 0.039 | Black-tailed deer | BC | Northwest | 1980-81 | Hatter 1988 |
| Yearling pregnancy | 0.83 | 0.108 | Mule deer | ID | Overall | 1998 | Hurley et al. 2011 |
| Yearling pregnancy | 0.92 | 0.07 | Mule deer | ID | Overall | 1999 | Hurley et al. 2011 |
| Yearling pregnancy | 0.625 | 0.17 | Mule deer | UT | Antimony | 1954-56 | Julander et al. 1961 |
| Yearling pregnancy | 0.86 | 0.12 | Mule deer | ID | Sublett | 1954-56 | Julander et al. 1961 |
| Yearling pregnancy | 0.11 | 0.098 | Mule deer | TX |  | 1990 | Lawrence et al. 2004 |
| Yearling pregnancy | 0.67 | 0.27 | Mule deer | TX |  | 1991 | Lawrence et al. 2004 |
| Yearling pregnancy | 0.68 | 0.1 | Mule deer | CA | Overall | 1997-2009 | Monteith et al. 2014 |
| Yearling pregnancy | 0.86 | 0.12 | Mule deer | SA |  | 2010 | Perera 2012 |
| Yearling pregnancy | 0.93 | 0.067 | Mule deer | SA |  | 2011 | Perera 2012 |
| Yearling pregnancy | 0.57 | 0.067 | Mule deer | UT |  | 1948-49 | Robinette and Gashwiler 1950 |
| Yearling pregnancy | 0.84 | 0.027 | Mule deer | UT |  | 1950-53 | Robinette et al. 1955 |
| Yearling pregnancy | 0.83 | 0.108 | Mule deer | UT |  | 1979 | Smith 1983 |
| Yearling pregnancy | 0.88 | 0.114 | Mule deer | UT |  | 1980 | Smith 1983 |
| Yearling pregnancy | 0.85 | 0.11 | Black-tailed deer | BC |  | 1963-67 | Thomas 1983 |
| Yearling pregnancy | 0.56 | 0.165 | Mule deer | MT |  | 1983-85 | Wood 1987 |

**CITATIONS**

Andelt, W. F., T. M. Pojar, and L. W. Johnson. 2004. Long‐term trends in mule deer pregnancy and fetal rates in Colorado. The Journal of Wildlife Management 68:542-549.

Bender, L. C., J. C. Boren, H. Halbritter, and S. Cox. 2011. Condition, survival, and productivity of mule deer in semiarid grassland-woodland in east-central New Mexico. Human-Wildlife Interactions 5:276-286.

Bender, L. C., and B. D. Hoenes. 2018. Age-related fecundity of free-ranging mule deer Odocoileus hemionus Cervidae in south-central, New Mexico, USA. Mammalia 82:124-132.

Bender, L. C., B. D. Hoenes, and C. L. Rodden. 2012. Factors influencing survival of desert mule deer in the greater San Andres Mountains, New Mexico. Human-Wildlife Interactions 6:245-260.

Bender, L. C., L. A. Lomas, and J. Browning. 2007. Condition, survival, and cause‐specific mortality of adult female mule deer in north‐central New Mexico. The Journal of Wildlife Management 71:1118-1124.

Bergman, E. J., C. J. Bishop, D. J. Freddy, G. C. White, and P. F. Doherty Jr. 2014. Habitat management influences overwinter survival of mule deer fawns in Colorado. The Journal of Wildlife Management 78:448-455.

Bishop, C. J., J. W. Unsworth, and E. O. Garton. 2005. Mule deer survival among adjacent populations in southwest Idaho. The Journal of Wildlife Management 69:311-321.

Bishop, C. J., G. C. White, D. J. Freddy, B. E. Watkins, and T. R. Stephenson. 2009. Effect of enhanced nutrition on mule deer population rate of change. Wildlife Monographs 172:1-28.

Bleich, V. C., B. M. Pierce, J. L. Jones, and R. T. Bowyer. 2006. Variance in survival of young mule deer in the Sierra Nevada, California. California Fish and Game 92:24.

Bleich, V. C., and T. J. Taylor. 1998. Survivorship and cause-specific mortality in five populations of mule deer. The Great Basin Naturalist:265-272.

Bowyer, R. T. 1991. Timing of parturition and lactation in southern mule deer. Journal of Mammalogy 72:138-145.

Bush, A. P. 2015. Mule deer demographics and parturition site selection: assessing responses to provision of water. University of Nevada, Reno.

Clark, J. S. 2022. Life History Trade-offs: The Effects of Habitat Selection on Columbian Black-tailed Deer Survival in Oregon.

DeCesare, N. J., C. Peterson, T. Hayes, C. Anton, D. Messmer, T. Chilton-Radandt, B. Lonner, E. Lula, T. Thier, N. Anderson, C. Loecker, C. Bishop, and M. Mitchell. 2021. Montana statewide mule deer study: ecology of mule deer in northern forests and integrated population modeling in the prairie-breaks. Final Report for Federal Aid in Wildlife Restoration Grant W-167-R. Montana Fish, Wildlife and Parks, Helena, Montana.

Dellinger, J., C. Shores, M. Marsh, M. Heithaus, W. Ripple, and A. Wirsing. 2018. Impacts of recolonizing gray wolves (Canis lupus) on survival and mortality in two sympatric ungulates. Canadian Journal of Zoology 96:760-768.

Farmer, C. J., D. K. Person, and R. T. Bowyer. 2006. Risk factors and mortality of black‐tailed deer in a managed forest landscape. The Journal of Wildlife Management 70:1403-1415.

Forrester, T. D., and H. U. Wittmer. 2019. Predator identity and forage availability affect predation risk of juvenile black‐tailed deer. Wildlife Biology 2019:1-12.

Freeman, E. D., R. T. Larsen, M. E. Peterson, C. R. Anderson Jr, K. R. Hersey, and B. R. Mcmillan. 2014. Effects of male‐biased harvest on mule deer: implications for rates of pregnancy, synchrony, and timing of parturition. Wildlife Society Bulletin 38:806-811.

Gilbert, S. L., K. J. Hundertmark, M. S. Lindberg, D. K. Person, and M. S. Boyce. 2020. The importance of environmental variability and transient population dynamics for a northern ungulate. Frontiers in Ecology and Evolution 8:531027.

Gogan, P. J., R. W. Klaver, and E. M. Olexa. 2019. Northern Yellowstone mule deer seasonal movement, habitat selection, and survival patterns. Western North American Naturalist 79:403-427.

Hall, J. T. 2018. Survival of Neonate Mule Deer Fawns in Southern Utah: Effects of Coyote Removal and Synchrony of Parturition. Brigham Young University.

Hamlin, K. L., S. J. Riley, D. Pyrah, A. R. Dood, and R. J. Mackie. 1984. Relationships among mule deer fawn mortality, coyotes, and alternate prey species during summer. The Journal of Wildlife Management:489-499.

Haskell, S. P., W. B. Ballard, J. T. Mcroberts, M. C. Wallace, P. R. Krausman, M. H. Humphrey, O. J. Alcumbrac, and D. A. Butler. 2017. Growth and mortality of sympatric white‐tailed and mule deer fawns. The Journal of Wildlife Management 81:1417-1429.

Hatter, I. 1988. Effects of wolf predation on recruitment of black-tailed deer on northeastern Vancouver Island. Wildlife Branch, Ministry of Environment.

Hatter, I. W., and D. W. Janz. 1994. Apparent demographic changes in black-tailed deer associated with wolf control on northern Vancouver Island. Canadian Journal of Zoology 72:878-884.

Hatter, I. W., and J. MacDermott. 2021. Demographic Changes and Fluctuations in a Black-tailed Deer (Odocoileus hemionus columbianus) Population on Northern Vancouver Island, British Columbia: Are They Related to Predation? Canadian Wildlife Biology & Management (CWBM) 10.

Heffelfinger, L. J., D. G. Hewitt, R. W. DeYoung, T. E. Fulbright, L. A. Harveson, W. C. Conway, and S. S. Gray. 2023. Shifting agriculture and a depleting aquifer: implications of row-crop farming on mule deer population performance. Animal Production Science.

Heffelfinger, L. J., K. M. Stewart, A. P. Bush, J. S. Sedinger, N. W. Darby, and V. C. Bleich. 2018. Timing of precipitation in an arid environment: Effects on population performance of a large herbivore. Ecology and Evolution 8:3354-3366.

Hellesto, R. A. 2023. Wheat Vs. Wild: Habitat Selection, Migration, and Parturition of Mule Deer in an Agricultural Landscape. Washington State University.

Hurley, M. A., M. Hebblewhite, P. M. Lukacs, J. J. Nowak, J. M. Gaillard, and C. Bonenfant. 2017. Regional‐scale models for predicting overwinter survival of juvenile ungulates. The Journal of Wildlife Management 81:364-378.

Hurley, M. A., J. W. Unsworth, P. Zager, M. Hebblewhite, E. O. Garton, D. M. Montgomery, J. R. Skalski, and C. L. Maycock. 2011. Demographic response of mule deer to experimental reduction of coyotes and mountain lions in southeastern Idaho. Wildlife Monographs 178:1-33.

Jackson, N. J., K. M. Stewart, M. J. Wisdom, D. A. Clark, and M. M. Rowland. 2021. Demographic performance of a large herbivore: effects of winter nutrition and weather. Ecosphere 12:e03328.

Johnstone-Yellin, T. L., L. A. Shipley, W. L. Myers, and H. S. Robinson. 2009. To twin or not to twin? Trade-offs in litter size and fawn survival in mule deer. Journal of Mammalogy 90:453-460.

Julander, O., W. L. Robinette, and D. A. Jones. 1961. Relation of summer range condition to mule deer herd productivity. The Journal of Wildlife Management 25:54-60.

Karish, T. 2022. Survival, activity patterns, movements, home ranges and resource selection of female mule deer and white-tailed deer in western Kansas. Kansas State University.

Kolar, J. L., J. J. Millspaugh, B. A. Stillings, C. P. Hansen, C. Chitwood, C. T. Rota, and B. P. Skelly. 2018. Potential effects of oil and gas energy development on mule deer in western North Dakota. North Dakota Game and Fish Department, Wildlife Division.

Lamb, S., B. R. McMillan, M. van de Kerk, P. B. Frandsen, K. R. Hersey, and R. Larsen. 2023. From conception to recruitment: Nutritional condition of the dam dictates the likelihood of success in a temperate ungulate. Frontiers in Ecology and Evolution 11:206.

Lawrence, R. K., S. Demarais, R. A. Relyea, S. P. Haskell, W. B. Ballard, and T. L. Clark. 2004. Desert mule deer survival in southwest Texas. The Journal of Wildlife Management 68:561-569.

Lindbloom, A., in Mule Deer Working Group, Western Association of Fish and Wildlife Agencies. 2021. 2021 Range-wide status black-tailed and mule deer.

Lomas, L. A., and L. C. Bender. 2007. Survival and cause‐specific mortality of neonatal mule deer fawns, north‐central New Mexico. The Journal of Wildlife Management 71:884-894.

Lukacs, P. M., G. C. White, B. E. Watkins, R. H. Kahn, B. A. Banulis, D. J. Finley, A. A. Holland, J. A. Martens, and J. Vayhinger. 2009. Separating components of variation in survival of mule deer in Colorado. The Journal of Wildlife Management 73:817-826.

Marescot, L., T. D. Forrester, D. S. Casady, and H. U. Wittmer. 2015. Using multistate capture–mark–recapture models to quantify effects of predation on age-specific survival and population growth in black-tailed deer. Population Ecology 57:185-197.

Matthews, P., and V. Coggins. Movements and mortality of mule deer in the Wallowa Mountains. 1997.

McCorquodale, S. M. 1999. Movements, survival, and mortality of black-tailed deer in the Klickitat Basin of Washington. The Journal of Wildlife Management:861-871.

McCoy, R., and S. Murphie. 2011. Factors affecting the survival of black-tailed deer fawns on the northwestern Olympic Peninsula, Washington. Makah Tribal Forestry Final Report, Neah Bay, Washington.

McCoy, R. H., S. L. Murphie, M. Szykman Gunther, and B. L. Murphie. 2014. Influence of hair loss syndrome on black‐tailed deer fawn survival. The Journal of Wildlife Management 78:1177-1188.

McNay, R. S., and J. M. Voller. 1995. Mortality causes and survival estimates for adult female Columbian black-tailed deer. The Journal of Wildlife Management:138-146.

Monteith, K. L., V. C. Bleich, T. R. Stephenson, B. M. Pierce, M. M. Conner, J. G. Kie, and R. T. Bowyer. 2014. Life‐history characteristics of mule deer: effects of nutrition in a variable environment. Wildlife Monographs 186:1-62.

Morano, S. 2016. Population ecology and summer habitat selection of mule deer in the White Mountains: implications of changing landscapes and variable climate. University of Nevada, Reno.

Nellis, C. H. 1964. Mule deer of the National Bison Range: population dynamics food habits and physical condition.

Nellis, C. H. 1968. Productivity of mule deer on the National Bison Range, Montana. The Journal of Wildlife Management:344-349.

Northrup, J. M., C. R. Anderson Jr, B. D. Gerber, and G. Wittemyer. 2021. Behavioral and demographic responses of mule deer to energy development on winter range. Wildlife Monographs 208:1-37.

Perera, A. 2012. Mule deer (Odocoileus hemionus) reproduction, fawn survival, and exposure of fawns to infectious agents in a chronic wasting disease endemic area of southern Saskatchewan. University of Saskatchewan.

Pojar, T. M., and D. C. Bowden. 2004. Neonatal mule deer fawn survival in west‐central Colorado. The Journal of Wildlife Management 68:550-560.

Quintana, N. T., W. B. Ballard, M. C. Wallace, P. R. Krausman, J. De Vos, O. Alcumbrac, C. Cariappa, and C. O'Brien. 2016. Survival of desert mule deer fawns in central Arizona. The Southwestern Naturalist 61:93-100.

Robinette, W. L., and J. S. Gashwiler. 1950. Breeding season, productivity, and fawning period of the mule deer in Utah. The Journal of Wildlife Management 14:457-469.

Robinette, W. L., J. S. Gashwiler, D. A. Jones, and H. S. Crane. 1955. Fertility of mule deer in Utah. The Journal of Wildlife Management 19:115-136.

Robinson, H. S., R. B. Wielgus, and J. C. Gwilliam. 2002. Cougar predation and population growth of sympatric mule deer and white-tailed deer. Canadian Journal of Zoology 80:556-568.

Schuyler, E. M., K. M. Dugger, and D. H. Jackson. 2019. Effects of distribution, behavior, and climate on mule deer survival. The Journal of Wildlife Management 83:89-99.

Shallow, J. R., M. A. Hurley, K. L. Monteith, and R. T. Bowyer. 2015. Cascading effects of habitat on maternal condition and life-history characteristics of neonatal mule deer. Journal of Mammalogy 96:194-205.

Skelton, N. K. 2010. Migration, dispersal, and survival patterns of mule deer (Odocoileus hemionus) in a chronic wasting disease-endemic area of southern Saskatchewan.

Smith, R. B. 1983. Mule deer reproduction and survival in the Lasal Mountains, Utah.

Sorensen, G. E. 2015. Ecology of adult female Rocky Mountain mule deer following habitat enhancements in North-Central New Mexico.

Speten, D. 2014. Assessment of mule deer fawn survival and birth site habitat attributes in south-central Oregon.

Symonds, K. K. 1992. Mule deer movements, survival, and use of contaminated areas at Rocky Flats, Colorado. Colorado State University.

Taber, R. D., and R. F. Dasmann. 1957. The dynamics of three natural populations of the deer Odocoileus hemionus columbianus. Ecology 38:233-246.

Thomas, D. C. 1983. Age-specific fertility of female Columbian black-tailed deer. The Journal of Wildlife Management 47:501-506.

Turnley, M. T., R. T. Larsen, T. A. Hughes, M. S. Hinton, D. W. Sallee, S. Lamb, K. R. Hersey, and B. R. McMillan. 2022. Are opportunistic captures of neonate ungulates biasing relative estimates of litter size? Animal Biotelemetry 10:38.

Unsworth, J. W., D. F. Pac, G. C. White, and R. M. Bartmann. 1999. Mule deer survival in Colorado, Idaho, and Montana. The Journal of Wildlife Management:315-326.

Washington Department of Fish and Wildlife. 2016. 2016 Game status and trend report. Wildlife Program, Washington Department of Fish and Wildlife, Olympia, Washington, USA.

Washington Department of Fish and Wildlife. 2021. 2021 Game status and trend report. Wildlife Program, Washington Department of Fish and Wildlife, Olympia, Washington, USA.

White, G. C., and R. M. Bartmann. 1998. Effect of density reduction on overwinter survival of free-ranging mule deer fawns. The Journal of Wildlife Management:214-225.

Wittmer, Heiko & Cristescu, Bogdan & Spitz, DB & Nisi, Anna & Wilmers, Christopher. (2021). Final Report Siskiyou Deer-Mountain Lion Study (2015-2020). 10.13140/RG.2.2.31758.28480.

Wood, A. K. 1987. Ecology of a prairie mule deer population. Montana State University-Bozeman, College of Letters & Science.

Wright, C. A., I. T. Adams, P. Stent, and A. T. Ford. 2020. Comparing Survival and Movements of Non‐Urban and Urban Translocated Mule Deer. The Journal of Wildlife Management 84:1457-1472.

Zager, P., G. Pauley, M. Hurley, and C. White. 2007. Statewide Ungulate Ecology. Idaho Dept. of Fish and Game, Boise, Idaho, USA.
