## Supporting Information S3 for "Variability and correlations among vital rates and their influence on population growth in mule and black-tailed deer"

*The Journal of Wildlife Management*

We noticed a discrepancy between studies reporting annual juvenile survival and those reporting summer and winter survival juvenile survival separately. Specifically, the product of the mean of the raw estimates of 0–6 month survival (*x̅* = 0.416, *N* = 37) and the mean of the raw estimates of 7–12 month survival (*x̅* = 0.588, *N* = 196) from our literature review imputes a 12 month survival rate of 0.244. This is substantially lower than the mean of the raw estimates of 12-month survival from studies that only presented an annual, first year survival rate (*x̅* = 0.307, *N* = 48). We have no explanation for this discrepancy, but we note that a similar difference was also present in the review by Forrester and Wittmer (2013).

Although our objective was to assess both summer and winter juvenile survival separately, and thus we could have discarded the studies that only presented 12-month juvenile survival, we decided to include them for two reasons. First, we wanted to have the largest sample size possible to most adequately characterize these vital rates, and second, in the event that the studies reporting 12-month survival were better representations of the true, underlying vital rate than were the product of the seasonal estimates, we did not want to introduce bias into our analyses and simulations.

We therefore estimated summer and winter juvenile survival from the studies reporting only 12-month survival estimates (*N* = 48). We did this by estimating the relative contributions of summer and winter survival toward annual survival, and decomposing each 12-month estimate into its constituent parts. Specifically, we realized that summer and winter survival rates could be estimated from annual survival rates using $S_{j sum}= {S_{j annual}}^{x\_sum}$ and $S_{j win}= {S_{j annual}}^{x\_win}$, where $x_{sum}= \frac{log(S_{j sum})}{log(S_{j sum \times}S_{j win})}$ and $x_{win}= \frac{log(S_{j win})}{log(S_{j sum \times}S_{j win})}$. We then used the posterior distributions for the resulting $S_{j sum}$ and $S_{j win}$ estimates as data in the subsequent logit-normal models. In this way we could both utilize all available data and ensure results weren’t biased by using a non-random subset of the data.

**LITERATURE CITED**

Forrester, T. D., and H. U. Wittmer. 2013. A review of the population dynamics of mule deer and black‐tailed deer O docoileus hemionus in North America. Mammal Review 43:292-308.
