## Supporting Information S4 for "Variability and correlations among vital rates and their influence on population growth in mule and black-tailed deer"

*The Journal of Wildlife Management*

**Table S1**: Estimates of the correlations (*ρ*) among pairs of mule and black-tailed deer vital rates fit using bivariate logit-normal hierarchical models summarized from a range-wide literature review. NA entries represent pairs of vital rates with insufficient data to fit models. Bold entries represent correlations whose 95% credible intervals do not overlap 0. *S* = survival, *P* = pregnancy, *LS* = litter size, and *j* = juvenile, *y* = yearling, and *a* = adult. $S_{j sum}$ = juvenile survival in summer (0–6 months) and $S_{j win}$ = juvenile winter survival (7–12 months).

| **Vital Rate 1** | **Vital Rate 2** | ***ρ*** | **95% CI** |
| --- | --- | --- | --- |
| $\boldsymbol{S}_{\boldsymbol{j sum}}$ | $\boldsymbol{S}_{\boldsymbol{j win}}$ | **-0.44** | **-0.63, -0.22** |
| $S_{j sum}$ | $S_{y}$ | -0.28 | -0.96, 0.79 |
| $S_{j sum}$ | $S_{a}$ | -0.11 | -0.48, 0.25 |
| $S_{j sum}$ | $P_{y}$ | 0.19 | -0.89, 0.96 |
| $S_{j sum}$ | $P_{a}$ | 0.57 | -0.36, 0.92 |
| $S_{j sum}$ | ${LS}_{y}$ | -0.14 | -0.92, 0.83 |
| $S_{j sum}$ | ${LS}_{a}$ | 0.26 | -0.07, 0.55 |
| $\boldsymbol{S}_{\boldsymbol{j win}}$ | $\boldsymbol{S}_{\boldsymbol{y}}$ | **0.79** | **0.02, 1.00** |
| $\boldsymbol{S}_{\boldsymbol{j win}}$ | $\boldsymbol{S}_{\boldsymbol{a}}$ | **0.23** | **0.01, 0.44** |
| $S_{j win}$ | $P_{y}$ | *NA* | *NA* |
| $S_{j win}$ | $P_{a}$ | -0.41 | -0.97, 0.75 |
| $S_{j win}$ | ${LS}_{y}$ | -0.33 | -0.97, 0.69 |
| $S_{j win}$ | ${LS}_{a}$ | 0.15 | -0.21, 0.5 |
| $S_{y}$ | $S_{a}$ | 0.56 | -0.04, 0.9 |
| $S_{y}$ | $P_{y}$ | *NA* | *NA* |
| $S_{y}$ | $P_{a}$ | *NA* | *NA* |
| $S_{y}$ | ${LS}_{y}$ | *NA* | *NA* |
| $S_{y}$ | ${LS}_{a}$ | *NA* | *NA* |
| $S_{a}$ | $P_{y}$ | 0.68 | -0.03, 0.98 |
| $\boldsymbol{S}_{\boldsymbol{a}}$ | $\boldsymbol{P}_{\boldsymbol{a}}$ | **0.59** | **0.15, 0.94** |
| $S_{a}$ | ${LS}_{y}$ | *NA* | *NA* |
| $S_{a}$ | ${LS}_{a}$ | 0.15 | -0.27, 0.56 |
| $P_{y}$ | $P_{a}$ | 0.55 | -0.24, 0.98 |
| $P_{y}$ | ${LS}_{y}$ | 0.20 | -0.6, 0.78 |
| $\boldsymbol{P}_{\boldsymbol{y}}$ | $\boldsymbol{LS}_{\boldsymbol{a}}$ | **0.70** | **0.17, 0.97** |
| $P_{a}$ | ${LS}_{y}$ | 0.44 | -0.83, 0.95 |
| $\boldsymbol{P}_{\boldsymbol{a}}$ | $\boldsymbol{LS}_{\boldsymbol{a}}$ | **0.24** | **0.04, 0.44** |
| $\boldsymbol{LS}_{\boldsymbol{y}}$ | $\boldsymbol{LS}_{\boldsymbol{a}}$ | **0.74** | **0.35, 0.92** |
