## Supporting Information S5 for "Variability and correlations among vital rates and their influence on population growth in mule and black-tailed deer"

**SUPPORTING INFORMATION S5: PREDICTIONS ABOUT LAMBDA AS A FUNCTION OF ADULT FEMALE SURVIVAL**

*The Journal of Wildlife Management*

**Table S2**: Predictions about lambda as a function of adult female survival rate in mule and black-tailed deer estimated using life-stage simulation analysis. Pr(λ ≥ 1) is the probability of observing a lambda of at least 1, indicating a stable or increasing population. The next columns provide predictions of lambda and 95% Bayesian Credible Intervals (BCIs).

| Adult Female Survival Rate (annual) | Pr(λ ≥ 1) | λ (mean) | λ (Lower 95% BCI) | λ (Upper 95% BCI) |
| --- | --- | --- | --- | --- |
| 0.60 | 0.00 | 0.74 | 0.63 | 0.88 |
| 0.65 | 0.00 | 0.79 | 0.68 | 0.94 |
| 0.70 | 0.01 | 0.84 | 0.73 | 0.99 |
| 0.75 | 0.10 | 0.89 | 0.78 | 1.04 |
| 0.80 | 0.27 | 0.95 | 0.84 | 1.09 |
| 0.85 | 0.52 | 1.01 | 0.89 | 1.15 |
| 0.90 | 0.73 | 1.06 | 0.95 | 1.21 |
| 0.95 | 0.94 | 1.12 | 0.99 | 1.27 |
| 1.0 | 1.00 | 1.17 | 1.05 | 1.33 |
